# TRPP Ion Channels Function in the *π*-Conformation Rather Than the *α*-Conformation

**DOI:** 10.64898/2026.11.04.764271

**Authors:** Chenfeng Zhang, Wojciech Kopec, Julien Michel

## Abstract

PKD2 (TRPP1) is a TRPP ion channel whose dysfunction causes autosomal dominant polycystic kidney disease (ADPKD). Understanding its conformational states and gating mechanism is important for elucidating its function and for structure-based drug discovery. Cryo-EM structures of WT-PKD2 contain a local *π*-helical turn in the S6 pore-lining helix (*π*-conformation). In contrast, a fully *α*-helical S6 (*α*-conformation) was observed in the gain-of-function mutant F604P-PKD2 and interpreted as the open state, leading to the proposed *π*-to-*α* activation mechanism. However, whether the *α*-conformation can support function has never been directly established. Here, using molecular dynamics simulations, we investigate the functional properties of the *α*- and *π*-conformations of PKD2. The *α*-conformation remains dewetted unless the lower gate is widened beyond the conformational range observed in mammalian TRP channels. By contrast, the *π*-conformation supports pore hydration and ion conduction following a small opening of the L677 gate, with simulated conductance comparable to experimental measurements. The closely related PKD2L1 (TRPP2) channel shows a similar dependence on conformation, with a hydrated *π*-conformation and a dewetted *α*-conformation. Together, these findings challenge the assignment of the *α*-conformation as the conductive state and support a model in which the *π*-conformation represents the functionally relevant state of TRPP channels. This model suggests that activation may involve an *α*-to-*π* transition rather than the previously proposed *π*-to-*α* transition, providing a new framework for understanding TRPP channel gating. This work sheds light on the molecular basis of polycystin channel dysfunction and could facilitate the development of PKD2-targeted therapeutic strategies for ADPKD.

**Significance statement:** Polycystin-2 (PKD2) is a TRPP-family ion channel whose dysfunction causes autosomal dominant polycystic kidney disease (ADPKD), one of the most common inherited kidney diseases. Two distinct conformations of TRPP channels have been identified by cryo-EM, termed the *α*- and *π*-conformations, and interpreted as different functional states, leading to the proposed *π*-to-*α* activation mechanism. Using molecular dynamics simulations, we challenge this mechanism and show that the *π*-conformation, rather than the *α*-conformation, is the functional state of PKD2 and the related channel PKD2L1. This revised view provides a new framework for understanding TRPP channel gating and guiding drug discovery for ADPKD. This work is also the first detailed atomistic simulation study of TRPP channel function, laying the groundwork for future computational research.

## Introduction

Autosomal dominant polycystic kidney disease (ADPKD) is the most common inherited kidney disease and is characterised by the progressive formation of renal cysts that can eventually lead to kidney failure.^1^ ADPKD is primarily caused by mutations in the PKD1 and PKD2 genes, which encode polycystin-1 (PC1) and polycystin-2 (PC2), respectively. PC1 and PC2 are components of two major channel complexes: the PC1-PC2 heterotetrameric channel, with a 1:3 stoichiometry, and the PC2 homotetrameric channel (hereafter referred to as PKD2). There is currently no cure for ADPKD. The approved drug tolvaptan can only slow disease progression and acts on downstream processes of the disease pathway rather than directly targeting PC1 or PC2.^2^ Therapeutic strategies that directly target the molecular defects underlying ADPKD are therefore of considerable interest. Recent industry efforts have begun to explore direct pharmacological modulation of polycystin ion-channel function,^3^ and the investigational drug VX-407, which targets specific PKD1 variants, is currently being evaluated in clinical trials.^4^ As PC2 is a key component of both major polycystin channel complexes, understanding the molecular basis of PKD2 function could provide broader insights into polycystin channel function and facilitate the development of PKD2-targeted therapeutic strategies.

PKD2, also known as TRPP1, belongs to the transient receptor potential (TRP) channel family, which is part of the voltage-gated ion channel (VGIC) superfamily. Like other TRP channels, PKD2 comprises a voltage-sensor-like domain (VSLD) formed by transmembrane helices S1–S4 and a pore domain (PD) formed by S5–S6. In addition, TRPP channels contain a characteristic extracellular domain between S1 and S2, known as the TOP domain (Tetragonal Opening for Polycystins, also called the polycystin domain)^5,6^ (Fig. 1A). Over the past decade, cryo-electron microscopy (cryo-EM) has produced a series of high-resolution structures of PKD2.^5–12^ With the exception of the gain-of-function (GOF) mutant F604P-PKD2, all other reported structures consistently show a *π*-bulge in the middle of the S6 pore-lining helix, characterised by a local distortion in which the backbone hydrogen-bonding pattern switches from *α*-helical (*i → i* + 4) to *π*-helical (*i → i* + 5). In contrast, structures of F604P-PKD2 show a standard *α*-helix in S6. The lower gate residue also differs between these conformations. While L677 occupies the constriction site in structures containing the *π*-bulge (*π*-conformation hereafter), F676 occupies the corresponding position in structures containing the standard *α*-helix (*α*-conformation hereafter) (Fig. 1C).

**Figure 1:**
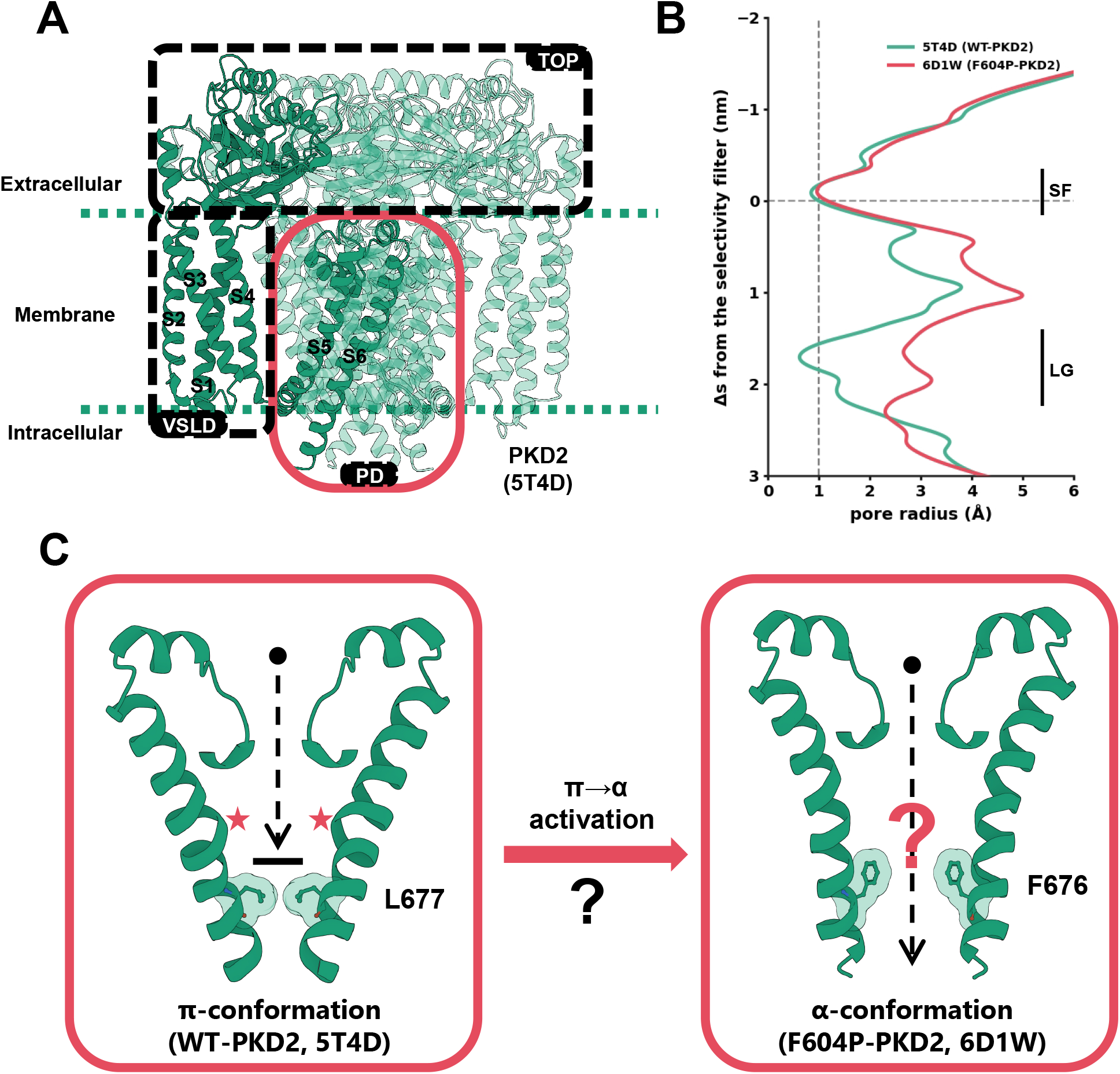
Structural basis of PKD2 and proposed *π*-to-*α* activation model. (**A**) Overall architecture of PKD2 (PDB 5T4D), showing the voltage-sensor-like domain (VSLD) formed by S1–S4, the TOP domain between S1 and S2, and the pore domain (PD) formed by S5–S6. (**B**) Pore radius profiles of PDB 5T4D (WT-PKD2) and PDB 6D1W (F604P-PKD2). The selectivity filter (SF) and lower gate (LG) regions are indicated. Grey dotted lines mark the SF position, which defines zero on the y-axis, and a pore radius of 1 Å. (**C**) Proposed *π*-to-*α* activation model and structures of the *π*- and *α*-conformations of the selectivity filter and the pore-lining S6 helix (residues 640–682). Two opposing subunits are shown for clarity. The *π*-conformation contains *π*-bulges (marked by red stars) and L677 at the lower gate, whereas the *α*-conformation adopts a standard *α*-helix with F676 at the lower gate. In the proposed model, the *π*-conformation is interpreted as a closed state that does not permit ion permeation, whereas the *α*-conformation is interpreted as an open state. However, there is no direct evidence that the *α*-conformation represents a functional state.

The first proposed gating mechanism of PKD2 suggested that the *π*-bulge acts as a hinge, allowing the bottom half of S6 to bend and open the lower gate.^6^ Another study proposed an activation mechanism involving the TOP domain and lipids.^7^ Both mechanisms were based on the assumption that PKD2 functions in the *π*-conformation. However, following the determination of the first cryo-EM structure of F604P-PKD2, the *α*-conformation was proposed to represent the open state of PKD2.^8^ This interpretation was mainly based on the pore radius observed in the structure (Fig. 1B). The pore radius of the F604P-PKD2 cryo-EM structure was considered sufficiently wide to permit ion permeation, whereas the pores of the *π*-conformations appeared to be in a closed state.^5–8^ In addition to these structural observations, functional data appeared to support this interpretation. F604P-PKD2, for which only *α*-conformation structures have been resolved, exhibits robust currents when expressed at the plasma membrane.^8,13^ In contrast, WT-PKD2, whose reported cryo-EM structures all adopt the *π*-conformation, has been reported to show no detectable plasma-membrane current.^6,8^ Together, these structural and functional observations were interpreted as supporting a model in which the *π*-conformation represents a closed state and the *α*- conformation an open state. This interpretation led to the proposal that PKD2 activation involves a *π*-to-*α* transition in S6^8^ (Fig. 1C), which was subsequently adopted by several studies.^11,14–16^

However, this interpretation remains an inference rather than a direct demonstration of the conductive state, and it relies on several assumptions that can be easily challenged. First, a pore that is geometrically wide enough to accommodate an ion does not necessarily support ion conduction, as the chemical properties of pore-lining residues can also affect the function of the channel. Second, cryo-EM structures capture conformational states under specific experimental conditions, which may not correspond to the conductive state of the channel. Third, the absence of detectable WT-PKD2 current at the plasma membrane does not necessarily indicate that the *π*-conformation is non-functional, as channel activity may depend on factors present in the native cellular environment but absent from the plasma membrane expression system. In fact, PKD2 has been reported to function in the primary cilium,^17,18^ one of its principal sites of cellular localisation.^19^ Finally, the behaviour of the GOF mutant F604P cannot be assumed to directly represent that of WT-PKD2, since a single mutation can substantially alter protein conformational dynamics and function.

These considerations lead to two important questions: does the *α*-conformation represent a functional state of WT-PKD2, and can the *π*-conformation instead represent a functional, conductive state? Answering these questions is important not only for understanding the gating mechanism of PKD2 but also for PKD2-targeted drug discovery. In particular, structure-based drug discovery relies on an understanding of structure-function relationships, making it important to establish the functional relevance of different PKD2 conformations. When assigning a conformation as a conductive state, molecular-level evidence that it can support ion conduction is therefore needed, rather than relying solely on its structural appearance and existing experimental data.

These questions are also directly relevant to the interpretation of previous experimental studies of PKD2. F604P-PKD2 has been widely used to investigate disease-associated mutations,^20^ small-molecule modulation, ^11,16^ protein-protein modulation,^21,22^ channel activation,^23^ and gating mechanisms.^8^ In some of these studies, the *α*-conformation has been interpreted as an activated or open state, and this structural assignment has been used to rationalise functional observations. Consequently, the interpretation of these observations depends, at least in part, on the functional state assigned to the underlying structure. Without direct molecular-level evidence, such assignments may lead to an incorrect understanding of PKD2 channel behaviour. Establishing whether the *α*- and *π*-conformations can support function therefore provides an important basis for interpreting these experimental observations.

Here, we use molecular dynamics simulations to assess the functional relevance of the *α*- and *π*-conformations of WT-PKD2. We first examine whether the *α*-conformation can support pore hydration and, if not, whether the required lower-gate opening falls within the range observed in mammalian TRP channel structures. We then investigate whether the *π*- conformation can support pore hydration and ion conduction following a controlled opening of the lower gate, and, if so, whether the resulting permeation properties are consistent with experimental observations. Finally, we extend the hydration analysis to PKD2L1 (TRPP2) to examine whether a similar relationship between S6 conformation and pore hydration is conserved across the TRPP family. Together, these analyses allow us to define the relationship between S6 conformation and channel function and to reassess the proposed functional role of the *α*- and *π*-conformations, as well as the proposed *π*-to-*α* activation mechanism.

## Results

### The *α*-conformation observed by cryo-EM does not support pore hydration

To investigate the functional properties of WT-PKD2 in the *α*-conformation, we first sought to establish a stable WT model adopting this conformation, as no cryo-EM structure of WT-PKD2 in the *α*-conformation has been reported. An initial WT-*α* model was therefore constructed by homology modelling using the experimentally determined *π*-conformation of WT-PKD2 as the template (Fig. S1A and Table S1, see Methods). This model underwent substantial structural relaxation, particularly in the S6 region, with one replica moving towards the *α*-conformation observed in the cryo-EM structure (Fig. S1B–D). Because this relaxation would consume a non-negligible fraction of the simulation time, we constructed a second WT-*α* model by directly reverting P604 to phenylalanine in the experimentally determined *α*-conformation structure of F604P-PKD2 (Fig. 2A). Across six independent 1 *µ*s replicas (Table S2), this model largely retained the initial conformation (Fig. S2A), although one subunit in one replica underwent a spontaneous transition towards the *π*-conformation (Fig. 2B). Together with the observation that all reported WT-PKD2 cryo-EM structures adopt the *π*-conformation, this suggests that WT-PKD2 may favour the *π*-conformation.

**Figure 2:**
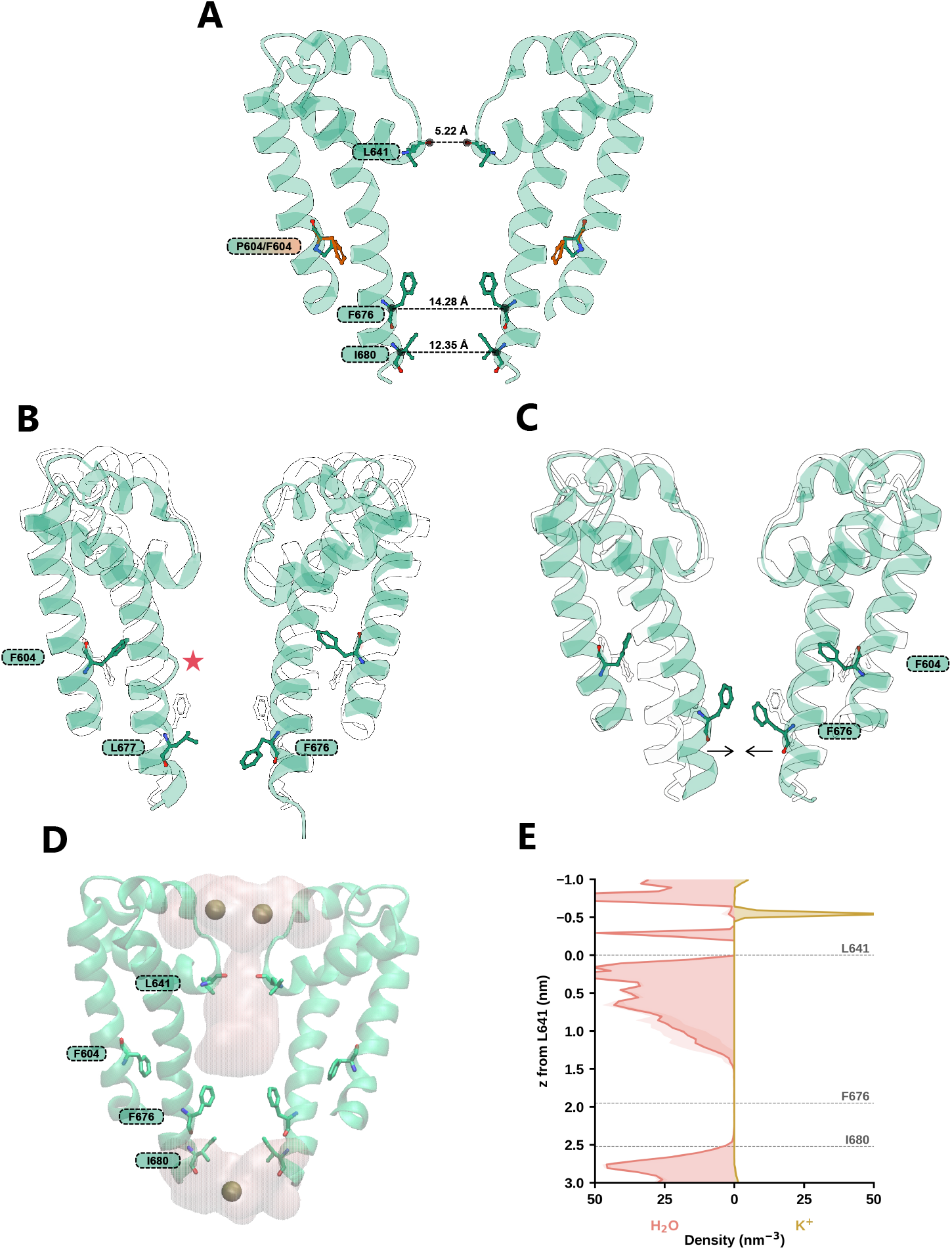
The *α*-conformation observed by cryo-EM does not support pore hydration. (**A**) WT *α*-conformation model based on PDB 6D1W (residues 598–682), with P604 (green) reverted to phenylalanine (orange). Two opposing subunits are shown for clarity. Inter-subunit distances are marked at the selectivity filter (L641 O) and the lower gate (F676 and I680 C*α*). (**B**) A spontaneous *α*-to-*π* transition during unbiased MD. The *π*-bulge is marked by a red star. Structures are superposed on the white initial model (A). (**C**) Spontaneous pore closure in one replica during unbiased MD. The final snapshot (green) of the replica is superposed on the white initial model (A). The F676 backbone moves towards the pore axis (arrows) and occludes the lower gate. (**D**) Representative snapshot of the *α*-conformation simulation with OPC water under an applied electric field, with the protein heavy atoms restrained to the initial model (A). Pore water is shown as a pink surface and K^+^ ions as spheres. (**E**) Water (left) and K^+^ (right) density within 3 Å of the pore axis in the simulations shown in (D), averaged over four 100-ns replicas (shading, SD across replicas). *z* is measured from the selectivity filter (L641 C*α*), with positive values towards the cytoplasm.

However, no continuous water pathway developed, and the pore spontaneously collapsed in all replicas. The S6 backbone moved towards the pore axis, while the F676 side chains rotated downward, further narrowing the pore (Figs. 2C and S2B). We also simulated the F604P-PKD2 model, which showed a similarly closed pore, indicating that pore collapse was not caused by reverting P604 to phenylalanine (Fig. S2C). Based on the difference between the residues modelled in the cryo-EM structure and the sequence of the construct used for cryo-EM, this pore collapse may be related to unresolved downstream helices that were not included in our model. Their absence may result in the loss of interactions that help stabilise the pore in the initial structure. Therefore, the observed pore collapse may not directly reflect the intrinsic stability of the *α*-conformation, and unbiased MD simulations alone were insufficient to assess its intrinsic functional properties.

To maintain the pore radius observed in the *α*-conformation cryo-EM structure (Fig. 1B), restraints were then applied to all heavy atoms of the protein to maintain the experimentally observed structure. Because the WT-*α* and F604P-*α* models were derived from the same cryo-EM structure, the restrained simulations preserve the same overall protein conformation. We therefore focused on WT-*α* for the restrained simulations. Multiple restrained simulations were performed, including simulations with an applied electric field to promote water entry into the pore. However, none of the simulations developed a continuous water pathway through the pore. Under an applied electric field, water occasionally bridged the lower gate between F676 and I680, but these events were transient, mostly lasting less than 0.5 ns and never exceeding 1.5 ns (Fig. S3A–D). We therefore considered whether the choice of water model could contribute to the observed dewetting, as different water models have been shown to affect channel behaviour.^24–26^ To test this possibility, we repeated the simulations using the OPC water model, which provides a more accurate description of the electrostatic and hydration properties of water. ^27^ Unexpectedly, the pore remained dewetted with OPC water (Fig. 2D,E). Inspection of the channel structure showed that the dewetted region was located between F676 C*ζ* and I680 C*α* (Fig. 2D). If this structure represents a functional conductive state, an ion would need to traverse an approximately 1 nm region (Fig. 2E) without a continuous hydration shell and coordinating residues, which would impose a substantial energetic penalty on ion permeation. Consistent with the MD simulations, a heuristic prediction using CHAP^28^ also showed that the pore would remain dewetted in the initial structure, with a contribution from F676 (Fig. S3E). Together, these results suggest that the *α*-conformation observed in the F604P-PKD2 cryo-EM structure is unlikely to represent a conductive open state.

### Pore hydration of the *α*-conformation requires gate opening beyond that observed in mammalian TRP channels or I680G mutation

Although the *α*-conformation cryo-EM structure lacks pore hydration, one could argue that it represents a closed state that could become hydrated upon pore opening. We therefore tested whether the *α*-conformation could support pore hydration when the lower gate was manually opened. F676 has been identified as a key lower-gate residue.^8,11^ We therefore scanned a series of diagonal C*α*–C*α* distance restraints between the four F676 residues, ranging from 14.5 to 19.0 Å in 0.5 Å increments (Figs. 3A and S4A). The corresponding distance in the cryo-EM structure is 14.28 Å (Fig. 2A). For each target distance, steered molecular dynamics was used to open the pore to the target distance over 10 ns, followed by a 100 ns production simulation with the corresponding C*α*–C*α* distance restraint. A continuous water pathway was first observed at a target distance of 18.5 Å (Fig. 3A). Notably, pore hydration occurred only during the final 5 ns of the production simulation at this target distance. Over the last 10 ns, the mean of the two diagonal C*α*–C*α* distances was 18.47 *±* 0.15 Å, fluctuating around the target distance. Interestingly, opening the F676 gate involved not only an increase in the C*α*–C*α* distance, but also a slight rotation of the F676 side chains away from the pore axis, while the L677 side chains moved slightly towards the pore axis (Fig. S5A). A *π*-bulge also formed during the simulation (Fig. 3B). These observations suggest that forced opening of the *α*-conformation can be accompanied by a structural rearrangement of S6 rather than simple dilation of the *α*-helical pore. In particular, these observations raise the possibility that pore opening may be accompanied by a transition towards the *π*-conformation.

**Figure 3:**
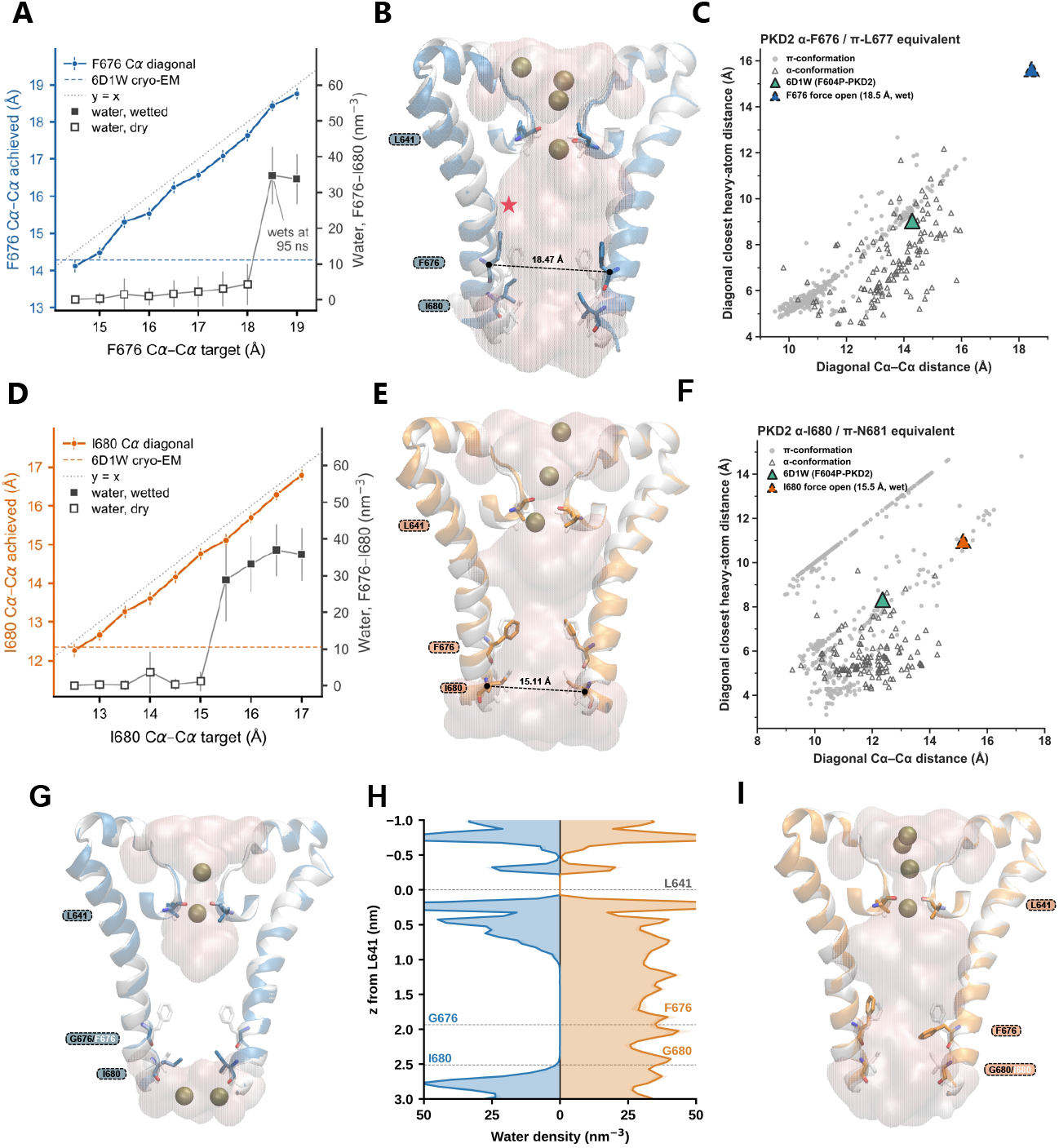
Pore hydration of the *α*-conformation requires gate opening beyond that observed in mammalian TRP channels or I680G mutation. **(A, D)** Forced opening of the pore by increasing the diagonal C*α*–C*α* distance between the four F676 (A) or I680 (D) residues. Water density was calculated between the F676 and I680 C*α* planes and within 3 Å of the pore axis. Mean SD values were calculated over the last 10 ns of each 100 ns production simulation. (**B, E**) Representative structures from the F676 (B) and I680 (E) distance scans at target diagonal C*α*–C*α* distances of 18.5 Å and 15.5 Å, respectively, at which the pore first becomes hydrated. The mean achieved distance over the last 10 ns is indicated at the corresponding residue. Widening at F676 (B) induces a *π*-bulge (red star). (**C, F**) Diagonal closest heavy-atom distance versus diagonal C*α*–C*α* distance between the F676-equivalent (C) and I680-equivalent (F) residues in mammalian TRP cryo-EM structures. The values for the hydrated *α*-conformation were calculated from the final frame of the first hydrated simulation. The hydrated *α*-conformation at F676 lies outside the range observed in the structures examined, while the hydrated *α*-conformation at I680 lies outside the range observed in the *α*-conformation structures. (**G, I**) Representative snapshots of the *α*-conformation simulation with F676G (G) or I680G (I) and the protein backbone atoms restrained to the initial model (Fig. 2A). I680G (I), but not F676G (G), results in pore hydration. (**H**) Water density for F676G (left) and I680G (right) within 3 Å of the pore axis, averaged over three 100-ns replicas (shading, SD across replicas). *z* is measured from the selectivity filter (L641 C*α*), with positive values towards the cytoplasm. All simulations were performed with OPC water under an applied electric field. For all snapshots, residues 640–682 are shown and structures are superposed on the white initial model (Fig. 2A). Pore water is shown as a pink surface and K^+^ ions as spheres.

As pore hydration can be achieved by forcing a pore sufficiently open, the observation of hydration alone does not establish that the required gate opening is physiologically accessible. We therefore compared the F676 geometry of the hydrated *α*-conformation with that of available cryo-EM structures of mammalian homotetrameric TRP channels (Table S3). For each structure, we measured the diagonal C*α*–C*α* distance and the diagonal closest heavy-atom distance between the F676-equivalent residues (Fig. 3C). Across the cryo-EM structures examined, the diagonal C*α*–C*α* distance did not exceed 16 Å, while the diagonal closest heavy-atom distance did not exceed 13 Å. In contrast, the hydrated *α*-conformation, taken from the final frame of the first hydrated simulation (target distance 18.5 Å), had a diagonal C*α*–C*α* distance of 18.44 Å and a diagonal closest heavy-atom distance of 15.56 Å. The hydrated *α*-conformation was therefore well outside the range observed in these mammalian homotetrameric TRP channel structures. These results indicate that hydration of the *α*- conformation is unlikely to occur at physiologically relevant gate geometries.

Further analysis of the pore radius during the F676 distance scan revealed that the pore was consistently narrower at I680 than at F676 (Fig. S5B). This also corresponds to the narrowest region of the lower gate in the cryo-EM structure (Fig. 1B). Notably, I680 has not been discussed as a potential gating residue in previous studies of F604P-PKD2.^8,11^ We therefore performed a second scan of the diagonal C*α*–C*α* distance between the four I680 residues using the same protocol, ranging from 12.5 to 17.0 Å in 0.5 Å increments, compared with 12.35 Å in the cryo-EM structure (Figs. 2A, 3D and S4A). A continuous water pathway emerged when the I680 distance reached approximately 15.5 Å (Figs. 3D,E). However, this opening was also substantially greater than that observed in available TRP *α*-conformation structures, where the corresponding C*α*–C*α* distance generally remained below 14.5 Å (Fig. 3F). Thus, as with F676, pore hydration required an opening beyond the conformational changes observed in available TRP channel structures. However, the I680 distance scan indicates that I680 is an important determinant of pore hydration in the *α*-conformation.

To further test the contribution of the lower-gate residues to pore hydration, we examined the effects of F676G and I680G in the *α*-conformation. In simulations with protein backbone atoms restrained to the initial model and an applied electric field, F676G did not result in pore hydration, suggesting that the dewetting observed in the *α*-conformation is not determined by F676 alone (Fig. 3G,H). In contrast, I680G resulted in the formation of a continuous water pathway under the same conditions (Fig. 3H,I). This result is consistent with the I680 distance scan and further supports an important role for I680 in pore hydration in the *α*-conformation.

To test the robustness of these results, we performed three additional sets of simulations (Table S2). First, to exclude a possible contribution of the selectivity filter to the dewetting, the distance scans were repeated with an additional lower-wall restraint on the distances between the L641 carbonyl oxygen atoms of the selectivity filter, which prevented the filter from closing (Fig. S4A). No pore hydration was observed in the F676 scan up to 19.0 Å, and hydration in the I680 scan required a larger opening of 16.0 Å (Figs. S6 and S7), indicating that keeping the selectivity filter open did not facilitate pore hydration and even hindered it. Second, because the downstream region of S6 is not resolved in the cryo-EM structure, we built an extended model of the *α*-conformation up to residue 694 and repeated both the distance scans and the mutation simulations. The results were similar to those of the original model, with pore hydration again requiring openings beyond the range observed in TRP channel structures or the I680G mutation (Figs. S8–S11). Third, to examine whether the water model affects these results, the distance scans and mutation simulations with the original model (up to residue 682) were repeated with the TIP3P water model. In these simulations, pore hydration was consistently less favourable with TIP3P than with OPC. No pore hydration was observed in the F676 scan up to 19.0 Å, and hydration in the I680 scan occurred only at a larger opening of 16.5 Å, compared with 15.5 Å with OPC (Figs. S12 and S13). Similarly, F676G did not result in pore hydration, whereas I680G showed only intermittent hydration of the lower gate (Fig. S14). Thus, the requirement for gate opening beyond the range observed in TRP channel structures, or for the I680G mutation, was not specific to the OPC water model.

Overall, these results show that the *α*-conformation is unlikely to represent a functional open state and identify I680 as an important contributor to the hydration barrier in the *α*-conformation.

### The *π*-conformation can open through subtle L677 movements and support experimental-level conductance

As the *α*-conformation is unlikely to represent a functional open state, we next asked whether the *π*-conformation could instead represent a functional state of PKD2. In the *π*-conformation, L677 forms the narrowest region of the pore (Fig. 1B). We first examined the role of L677 by introducing the L677G mutation (Fig. 4A,B). In unbiased MD simulations, the region between residues 677 and 681 became hydrated, but the pore below the selectivity filter dewetted at F669 and L673 owing to spontaneous pore closure (Fig. S15). When the protein backbone atoms were restrained to the initial model under an applied electric field, a continuous water pathway formed, although a K^+^ ion blocked the pore at N681 (Fig. 4B,C). This result suggests that L677 plays a key role in controlling pore hydration in the *π*-conformation.

**Figure 4:**
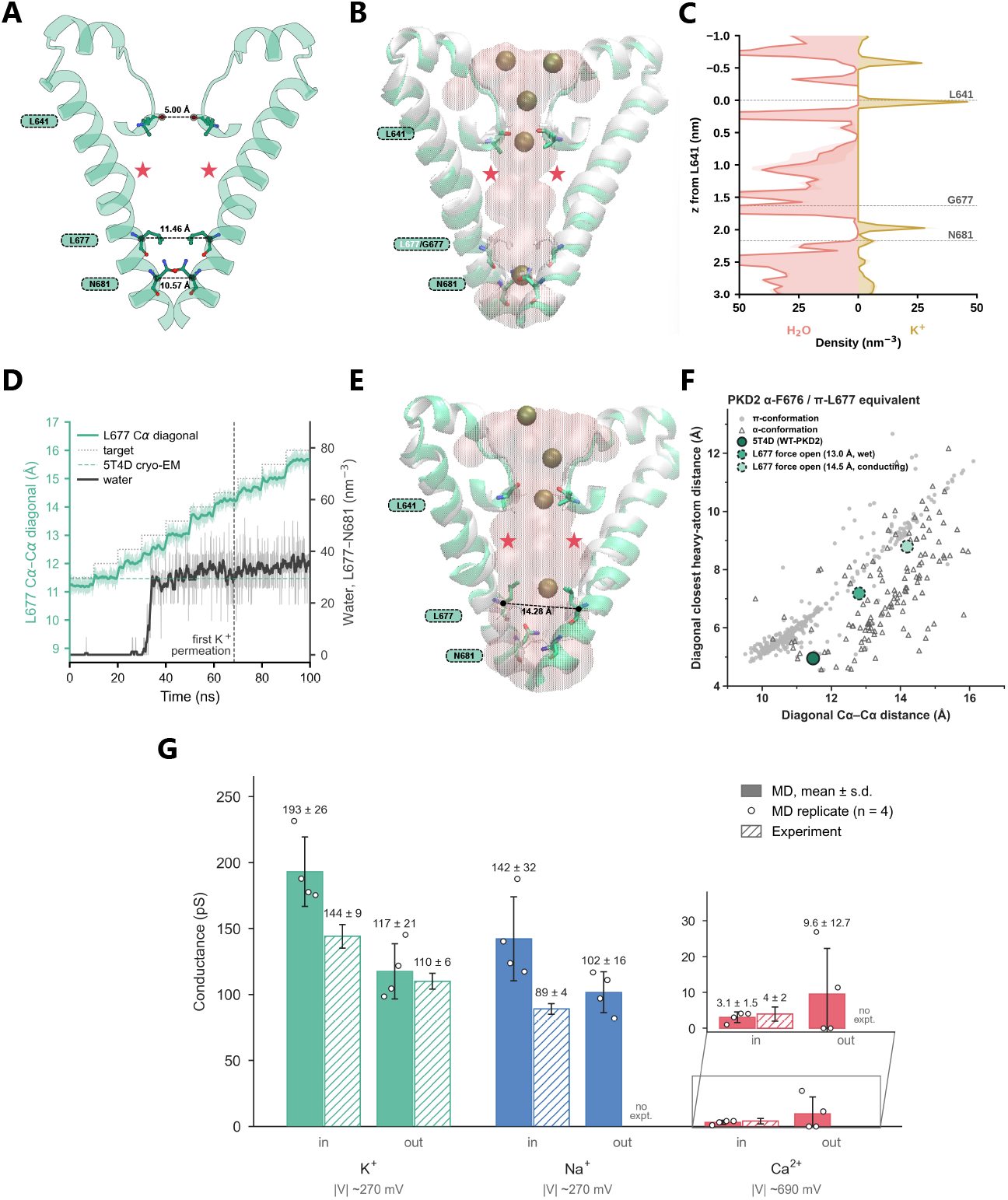
The *π*-conformation can open through subtle L677 movements and support experimental-level conductance. (**A**) *π*-conformation model (PDB 5T4D, residues 640–682). Two opposing subunits are shown for clarity. Inter-subunit distances are marked at the selectivity filter (L641 O) and the lower gate (L677 and N681 C*α*). (**B**) Representative snapshot showing pore hydration of the L677G mutant with the protein backbone atoms restrained to the white initial model (A). (**C**) Water (left) and K^+^ (right) density within 3 Å of the pore axis, averaged over three 100-ns replicas (shading, SD across replicas). *z* is measured from the selectivity filter (L641 C*α*), with positive values towards the cytoplasm. (**D**) Stepwise increase of the diagonal C*α*–C*α* distance between the four L677 residues. The distance was increased by 0.5 Å every 10 ns. Water density was calculated between the L677 and N681 C*α* planes and within 3 Å of the pore axis. Thin lines show the raw data and thick lines show the 1-ns running mean. (**E**) Representative snapshot from the L677 distance scan at a target diagonal C*α*–C*α* distance of 14.5 Å. The achieved distance is indicated. (**F**) Diagonal closest heavy-atom distance versus diagonal C*α*–C*α* distance between the L677-equivalent residues in mammalian TRP cryo-EM structures. The open *π*-conformation falls within the range observed in the structures. (**G**) Simulated conductance of K^+^, Na^+^, and Ca^2+^ for WT-PKD2 in the *π*-conformation, with the lower gate held open by restraining the diagonal L677 C*α*–C*α* distance to 14.5 Å. Values are mean SD over four replicas, simulated with CHARMM36m/TIP3P and ECC (see Methods). *V* is the applied trans-membrane potential. Experimental values are from single-channel recordings of WT-PKD2 in native primary cilia. ^18,29^ Simulations in (B)–(F) were performed with OPC water under an applied electric field. For all snapshots, *π*-bulges are marked by red stars, pore water is shown as a pink surface and K^+^ ions as spheres.

To further investigate whether the *π*-conformation could support pore opening, we performed a similar distance scan to that used for the *α*-conformation, this time varying the diagonal C*α*–C*α* distance between the four L677 residues. The L677 side chains were restrained to adopt an open-like rotamer from PDB 5MKE (Figs. 4E and S4B). The L677 distance was varied from 11.5 to 16.0 Å in 0.5 Å increments, compared with 11.48 Å in the cryo-EM structure (Fig. 4A,D). Unlike in the *α*-conformation, a continuous water pathway developed even during a rapid scan in which each distance was maintained for only 10 ns within a single 100 ns simulation, suggesting a substantially lower barrier to pore hydration (Fig. 4D). The pore became hydrated when the L677 distance reached approximately 12.5 Å (target distance 13 Å), and full ion permeation was observed when the distance reached 14 Å (target distance 14.5 Å, Fig. 4D,E). Increasing the L677 distance to a target of 14.5 Å required each L677 residue to move by only approximately 1.5 Å from its initial position. Such a small conformational change may be difficult to resolve at the nominal 3 Å resolution of the cryo-EM structure. Moreover, unlike the *α*-conformation, this *π*-open conformation falls within the range of conformations observed in available TRP structures (Fig. 4F). Even without the open-rotamer restraint on L677, a continuous water pathway and ion permeation were still observed at a wider opening that remained within the range of TRP channel structures (Fig. S16). These results suggest that the *π*-conformation can more readily support pore opening and can become conductive through relatively small movements of the L677 lower gate.

To further characterise the ion-conducting ability of the *π*-conformation, we then tested ion permeation at an L677 diagonal C*α*–C*α* distance of 14.5 Å. Using the AMBER ff19SB force field with the OPC water model, ion permeation was substantially underestimated (Fig. S17A). This is consistent with the known limitations of conventional fixed-charge force fields, which do not explicitly account for electronic polarisation and can therefore over-estimate ion–residue interactions. ^30,31^ To address this limitation, we applied the electronic continuum correction (ECC) to obtain a more accurate estimate of ion conductance. We transferred the WT-*π* system back to the CHARMM force field with the TIP3P water model for the ECC simulations, as the water pathway in the *π*-conformation was not affected by the choice of TIP3P or OPC water models (Fig. S18), and ECC is more established for the CHARMM force field.^32,33^ The resulting conductance values for K^+^, Na^+^, and Ca^2+^ were comparable in magnitude to the available experimental values (Fig. 4G), and the relative conductance followed the experimental trend with K^+^ *>* Na^+^ *≫* Ca^2+^.^6,18,29^ We further simulated outward conduction (Fig. 4G). For K^+^, the only ion for which outward single-channel data are available, the outward conductance was lower than the inward conductance, consistent with experimental observations.^29^

The differences in absolute conductance between simulation and experiment arise mainly from two factors. First, the ECC charge-scaling factors were adopted from studies of other channels rather than optimised for PKD2.^32,33^ The conductance was sensitive to this choice, as using the theoretical scaling factor of 0.75^31^ instead of 0.78 led to a higher K^+^ conductance (Fig. S17B). Second, the 14.5 Å L677 opening may be larger than required for ion permeation, as a continuous water pathway was already established at approximately 12.5 Å. However, our primary aim was to determine whether the *π*-conformation could support ion conduction at an experimentally relevant level, and the differences in the absolute conductance values do not preclude this conclusion. Overall, these results support the *π*-conformation as a conductive state of WT-PKD2.

### PKD2L1 shows similar conformational dependence of pore hydration

PKD2L1 (TRPP2), a close family member of PKD2, has also been proposed to adopt an open state in the *α*-conformation,^34^ while the *π*-conformation modelled from PKD2 has been treated as a closed state.^35–37^ To examine whether a similar relationship between conformation and pore hydration exists in PKD2L1, we performed simulations of its *α*- and *π*-conformations (Fig. 5A,C). Both models were positioned near the edge of the conformational range defined by the equivalent-residue plot (Fig. 5B), indicating that both models are in a relatively open state compared with other TRP channel structures. In the *α*- conformation, the pore collapsed during unbiased MD simulations (Fig. S19A), and even when the protein heavy atoms were restrained to the initial model under an applied electric field, no continuous water pathway formed (Fig. 5D,E). This dewetting is consistent with a CHAP heuristic prediction (Fig. S19B). Although its pore is wider than that of the PKD2 *α*- conformation model (Figs. 2A and 5A), the hydrated region extended only to F556 (F676 in PKD2), and the pore remained blocked by I560 (Fig. 5D,E). This is consistent with the role of the corresponding I680 residue in limiting pore hydration in the *α*-conformation of PKD2. In contrast, the *π*-conformation formed a continuous water pathway with all protein heavy atoms restrained to the initial model (Fig. 5E,F). Surprisingly, a continuous water pathway also formed spontaneously during unbiased MD simulations, even without an applied electric field (Fig. S20A,B). Although the model was still relaxing during these simulations, this relaxation did not proceed towards the *α*-conformation. In addition, A558 moved towards a pore-facing orientation in two adjacent subunits in each of the three replicas, consistent with the proposed L557–A558 dual gate in PKD2L1^8^ (Fig. S20C,D). Together, these results show that the *π*-conformation of PKD2L1 supports pore hydration, whereas the *α*-conformation does not. This behaviour is consistent with our observations for PKD2 and suggests that the relationship between the *π*-conformation and pore hydration may extend beyond PKD2 to other members of the TRPP family.

**Figure 5:**
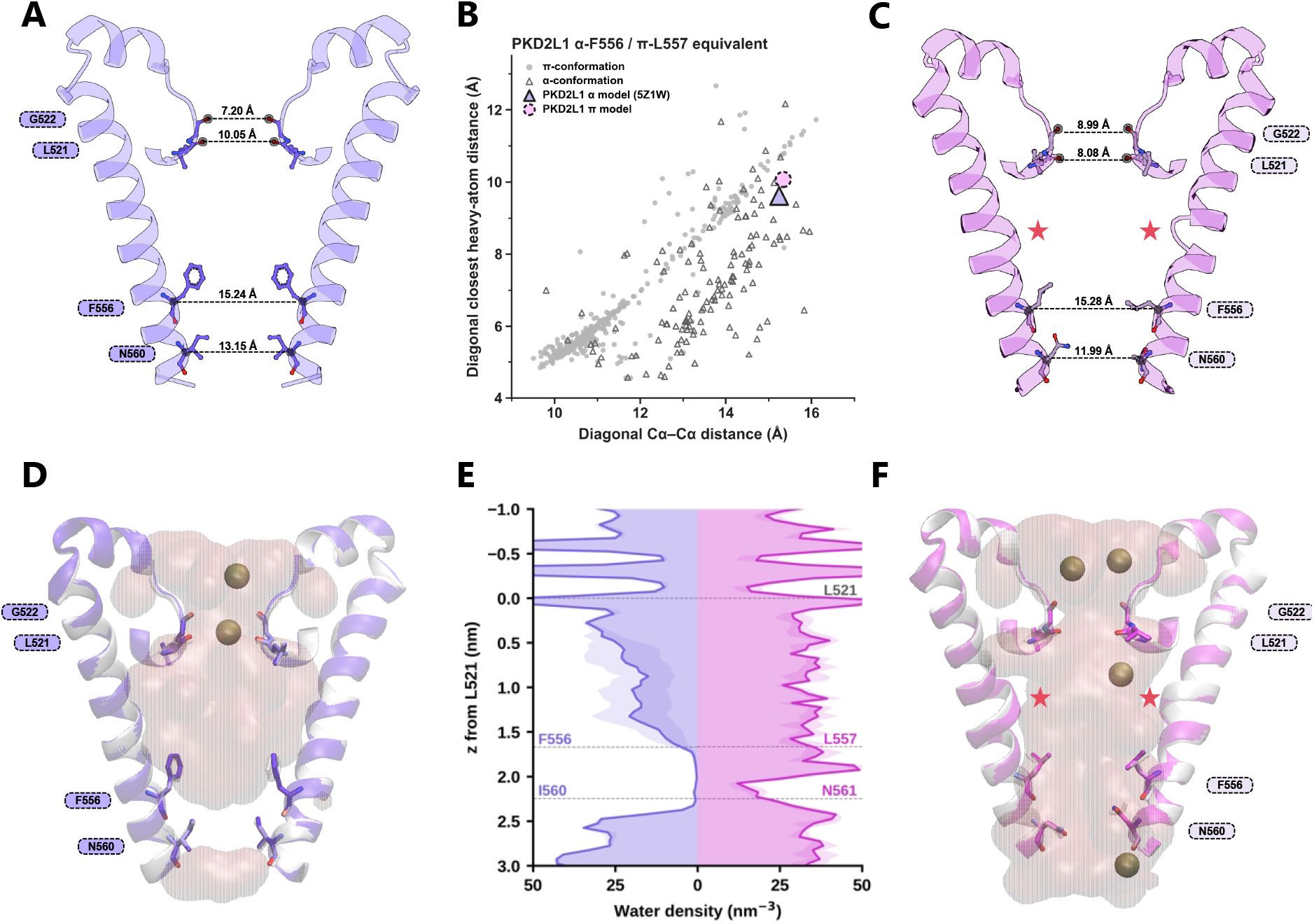
PKD2L1 shows similar conformational dependence of pore hydration. (**A,C**) *α*-conformation (A) and *π*-conformation (C) models of PKD2L1 (residues 520–563). The *α*-conformation model was obtained with Boltz-2 using PDB 5Z1W as a template, and the *π*-conformation model was constructed with MODELLER from the *α*-conformation model (see Methods). Two opposing subunits are shown for clarity. Inter-subunit distances are marked at the selectivity filter (L521 O) and the lower gate (F556 and I560 C*α* for the *α*-conformation, and L557 and N561 C*α* for the *π*-conformation). (**B**) Diagonal closest heavy-atom distance versus diagonal C*α*–C*α* distance between the *α*-F556/*π*-L557-equivalent residues of PKD2L1 (corresponding to *α*-F676/*π*-L677 in PKD2) in mammalian TRP cryo-EM structures. The PKD2L1 *α*- and *π*-conformations are positioned near the edge of the conformational range. (**D**) Representative snapshot from the restrained *α*-conformation simulation of PKD2L1, with the protein heavy atoms restrained to the white initial model (A) under an applied electric field. No continuous water pathway formed. (**E**) Water density within 3 Å of the pore axis, in the restrained *α*-conformation (left) and *π*-conformation (right) simulations, averaged over four 100-ns replicas (shading, SD across replicas). *z* is measured from the selectivity filter (L521 C*α*), with positive values towards the cytoplasm. (**F**) Representative snapshot from the restrained *π*-conformation simulation of PKD2L1, with the protein heavy atoms restrained to the white initial model (C) under an applied electric field. A continuous water pathway formed. All simulations were performed with OPC water. For all snapshots, *π*-bulges are marked by red stars, pore water is shown as a pink surface and K^+^ ions as spheres.

## Discussion

Since the structures of F604P-PKD2 and PKD2L1 were reported, the *α*-conformation has been widely interpreted as the open state of TRPP channels, leading to the proposal that activation involves a *π*-to-*α* transition.^8,34^ Here, we show that the *α*-conformation observed in the F604P-PKD2 cryo-EM structure, which has been interpreted as an open state, does not support a continuous water pathway for ion permeation. Pore hydration in the *α*- conformation requires either opening the lower gate beyond the conformational range observed in mammalian homotetrameric TRP channels or mutating I680 to glycine. In contrast, the *π*-conformation can support experimentally relevant ion conductance with a small opening of the L677 gate that lies within the range of TRP channel structures and below the resolution of the cryo-EM structure. We further extend this conformation–function relationship to PKD2L1 (TRPP2), showing that its *π*-conformation readily supports pore hydration, whereas its *α*-conformation remains dewetted. These findings challenge the assignment of the *α*-conformation as the functional state and instead support the *π*-conformation as the functional state of TRPP channels (Fig. 6A). This further suggests that activation may involve an *α*-to-*π* transition rather than the previously proposed *π*-to-*α* transition (Fig. 6A). The assignment of functional states from cryo-EM structures often relies on pore geometry, with a sufficiently large pore radius taken as an indication that ions can pass through. PKD2 and PKD2L1 illustrates the limitations of this approach. Although the pore radius of the *α*-conformation is geometrically sufficient to accommodate a bare ion, our simulations show that the cryo-EM structure does not support the continuous pore hydration required for ion permeation. Similar limitations have been identified in other ion channels, where open-state assignments based primarily on pore geometry have been challenged by molecular dynamics simulations.^38–40^ These findings highlight that pore geometry alone is insufficient to establish whether a structure represents a functional conductive state, and that the chemical environment within the pore must also be considered. Heuristic approaches that account for pore hydrophobicity alongside geometry can help identify potential barriers to pore hydration and provide a more informed assessment of the conductive potential of cryo-EM structures.^41^

**Figure 6:**
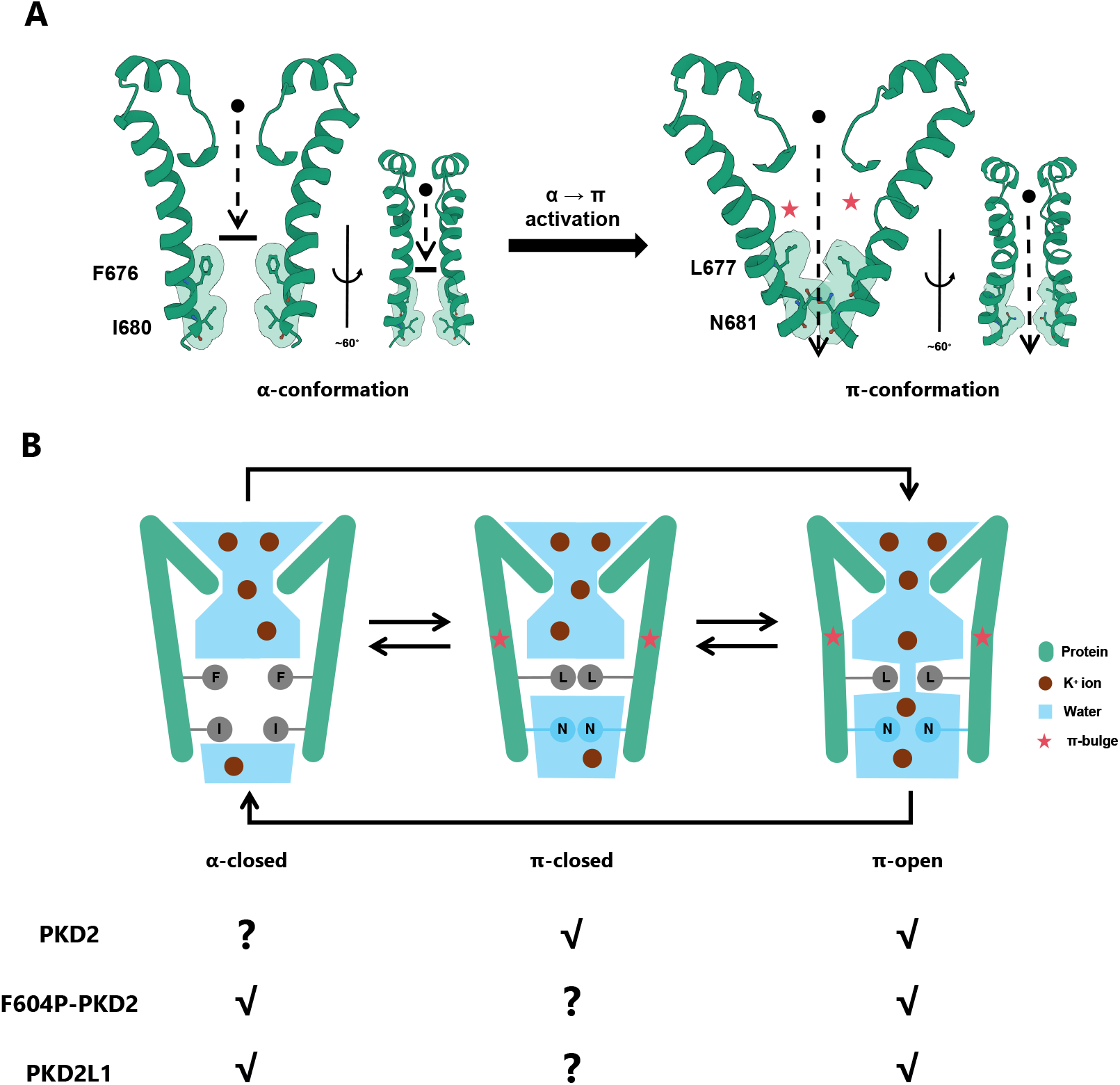
Proposed model of TRPP channel gating. (**A**) Structures of the *α*- and *π*-conformations of the selectivity filter and the pore-lining S6 helix (residues 640–682) of PKD2, illustrating the proposed *α*-to-*π* activation. The *α*-conformation is taken from PDB 6D1W and the *π*-conformation from the final frame of an ion permeation simulation with the gate opened. Two opposing subunits are shown for clarity, in two views rotated by approximately 60*^◦^*. The *α*-conformation does not permit ion permeation, whereas the *π*- conformation does. Key lower-gate residues are shown as sticks with a transparent surface. *π*-bulges are marked by red stars. (**B**) Proposed three-state model of TRPP channels. In the *α*-closed state, the pore is not physically occluded, but the hydrophobic lower-gate residues phenylalanine (F) and isoleucine (I) create a vapour-like region that prevents water and ion permeation, consistent with hydrophobic gating. In the *π*-closed state, leucine (L) physically occludes the pore and prevents water and ion permeation, whereas the hydrophilic asparagine (N) below it allows water entry. In the *π*-open state, a small opening at leucine allows water and ion permeation. The *π*-closed and *π*-open states are consistent with a helix-bundle-crossing gate. Arrows indicate potential transitions between states. The table summarises the states accessible to WT-PKD2, F604P-PKD2 and PKD2L1 based on experimental and simulation data (✔) and those that remain undetermined (?). F, I, L and N correspond to F676, I680, L677 and N681 in PKD2 and F556, I560, L557 and N561 in PKD2L1.

The different hydration behaviours of the *α*- and *π*-conformations suggest that hydrophobic and steric effects may contribute differently to gating in the two states. In the *α*- conformation, the pore remained dewetted even in the absence of a geometric constriction, suggesting that the dewetting is not mainly due to a steric barrier (Fig. 6B). This behaviour is consistent with hydrophobic gating, in which the chemical properties of the pore contribute to the free-energy barrier to hydration.^42,43^ In particular, the lower gate contains the hydrophobic residues F676 and I680, with our simulations identifying I680 as an important determinant of pore hydration. In contrast, the *π*-conformation more closely resembles a helix-bundle-crossing gate, in which a small opening at L677 that removes the steric obstruction is enough for pore hydration (Fig. 6B). Although L677 is hydrophobic, the hydrophilic residue N681 one helical turn below it may contribute to the greater ease of pore wetting in this conformation. Interestingly, a similar pattern is observed across the mammalian TRP channel structures examined in this study (Table 1). With the exception of TRPA1, the *α*-conformations generally contain two hydrophobic residues at the lower gate, whereas the *π*-conformations contain one hydrophobic residue together with either a hydrophilic or a small residue. This conserved pattern suggests that the relationship between S6 conformation and pore hydration may extend beyond TRPP channels to the broader TRP family.

**Table 1:** Pore-lining residue pairs measured in each channel. Each entry gives the two residues measured, one helical turn apart, as one-letter codes. They are the positions equivalent to PKD2 L677 and N681 in the *π*-conformation and to F676 and I680 in the *α*-conformation. For TRPM8 in the *α*-conformation, F979 and V983 were used, chosen by visual inspection. Detailed residue numbering is given in Table S4.

| **Channel** | **π** | **α** | **Channel** | **π** | **α** |
|---|---|---|---|---|---|
| **TRPA** |  |  | **TRPP** |  |  |
| TRPA1 | I+V | L+A | PKD2 | L+N | F+I |
|  |  |  | PKD2L1 | L+N | F+I |
| **TRPC** |  |  | **TRPML** |  |  |
| TRPC3 | I+N | L+I | TRPML1 | I+T | F+I |
| TRPC4 | I+N | L+M | TRPML2 | I+T | F+I |
| TRPC5 | I+N | L+M | TRPML3 | I+T | F+I |
| TRPC6 | I+N | L+I |  |  |  |
| **TRPM** |  |  | **TRPV** |  |  |
| TRPM2 | I+N | L+F | TRPV1 | I+G | L+M |
| TRPM3 | I+N | L+F | TRPV2 | I+S | L+M |
| TRPM4 | I+S | L+F | TRPV3 | I+G | L+M |
| TRPM5 | I+S | L+F | TRPV4 | I+G | L+M |
| TRPM7 | I+N | L+F | TRPV5 | I+G | F+M |
| TRPM8 | V+G | F+V | TRPV6 | I+G | L+M |

Our simulations provide several lines of evidence suggesting that the *α*-conformation can transition towards the *π*-conformation in WT-PKD2. These include a spontaneous conformational transition from *α*- towards the *π*-conformation in unbiased MD simulations and the observation that large pore opening is associated with formation of the *π*-bulge and rotation of the side chains at the lower gate. Together with the distinct functional properties of the *α*- and *π*-conformations observed in PKD2 and PKD2L1, these observations support a model in which TRPP channels may undergo *α*-to-*π* activation rather than the previously proposed *π*-to-*α* activation. This proposed mechanism would place TRPP channels within the broader framework of TRP channel gating.^14^ Across the TRP family, the functional conductive states of most channels have been characterised in the *π*-conformation, whereas the closed state can adopt either the *π*- or *α*-conformation.^14^ Here, we propose that TRPP channels can access *α*-closed, *π*-closed, and *π*-open states, similar to the proposed state model for TRPV channels (Fig. 6B).^40^ Based on experimental observations of TRPP channel behaviour at the plasma membrane and in the primary cilium, we suggest that WT-PKD2 predominantly occupies the *π*-closed state at the plasma membrane, while factors present in the primary cilium facilitate access to the *π*-open state. For the F604P mutant, the *α*-closed state is introduced, while the *π*-open state becomes accessible at the plasma membrane. PKD2L1, unlike PKD2, can access both the *α*-closed and *π*-open states at the plasma membrane and may also access a *π*-closed state.

The identification of the *π*-conformation as a functional state and the proposed *α*-to-*π* activation model do not conflict with previous experimental observations that were interpreted with the *α*-conformation as the functional state. Instead, they may provide a more consistent explanation for these observations. For example, in the study that identified gating residues in PKD2 and PKD2L1,^8^ F676 and L677 were proposed to form a double-residue gate based on mutational data from F604P-PKD2. The mutation data showed that F676A/G had no effect on current, whereas F676Y/W reduced current. L677G increased current, whereas the larger hydrophobic substitutions L677F/Y/W reduced current. F676 was considered a gating residue based on its orientation towards the pore in the *α*-conformation, and the reduced current caused by F676Y/W further supported this interpretation, as the larger side chains were proposed to increase pore blockage. Unlike F676, L677 was considered a gating residue based purely on the relationship between the size of the substituted residue and the resulting current. However, this double-residue gate interpretation cannot explain the lack of effect of F676A/G and how F676 and L677 could function together if the *α*-conformation is the functional state. Moreover, our simulations showed that L677G did not promote pore hydration in the *α*-conformation (Fig. S21), suggesting that the increased current observed for L677G is unlikely to arise from the *α*-conformation. By contrast, if the *π*-conformation is considered the functional state, these data can be better explained. First, the effects of L677 mutations are consistent with L677 forming the lower gate in the functional *π*-conformation. Second, the effects of F676 substitutions can be related to the *α*-to-*π* transition, as F676 points towards the membrane in the *π*-conformation and therefore needs to rotate away from the pore during the transition. The larger Y/W substitutions may hinder this rotation, whereas the smaller A/G substitutions may have little effect, providing a possible explanation for their different effects on channel current. Beyond F604P-PKD2, the gating residues identified in WT-PKD2 (L677) and PKD2L1 (L557 and A558)^8^ are consistent with our simulation results supporting the *π*-conformation as the functional state. Together, our proposed model and activation framework are compatible with the available experimental data while providing a more consistent explanation for previous experimental findings.

Several limitations should be considered when interpreting these findings. First, although the OPC water model provides a more realistic description of water, it remains a classical model and may not fully reproduce the behaviour of water in a confined pore. To our knowledge, there is currently no evidence that OPC shows anomalous behaviour in protein pores, but this possibility cannot be completely excluded. Second, the WT-*α* model was generated by reverting P604 to phenylalanine in the F604P-PKD2 structure rather than from an experimentally resolved structure, and may therefore differ from the native WT-*α* conformation. As the pore region relevant to our analysis is conserved, such differences are unlikely to alter our conclusions, although they may affect quantitative details. Third, the open state of the *π*-conformation was generated with restraints and has not been resolved experimentally. Nevertheless, it is likely to represent a realistic conductive state, as it required only small movements of L677 that are within the resolution of the cryo-EM structure, lies within the range of TRP channel structures and reproduces experimental-level conductance. Finally, this study focuses on how the *α*- and *π*-conformations of the pore relate to channel function rather than on the complete gating process of PKD2 or PKD2L1. Conformational changes in other regions of the channel and their coupling to the pore were not examined, and preliminary free-energy calculations of the S6 *α*–*π* transition were difficult to converge (Fig. S22). Even a converged free-energy landscape would reflect only the conformational preference under the simulated conditions and would not by itself establish which conformation is functional. Future studies that directly characterise the *α*-to-*π* transition and its coupling to the rest of the channel will therefore be important for evaluating the proposed activation mechanism.

Overall, our findings challenge the conventional view that the *α*-conformation represents the open state and instead support the *π*-conformation as the functional state of TRPP channels. The proposed *π*-to-*α* activation mechanism should therefore be reconsidered. We propose a revised framework in which TRPP channels can access distinct *α*-closed, *π*-closed, and *π*-open states, with activation involving transitions towards the functional *π*-open state. This framework provides a new basis for understanding TRPP channel gating and for identifying functionally relevant conformations for structure-based PKD2-targeted drug discovery.

## Materials and Methods

### System preparation

All models were built based on three deposited cryo-EM structures: wild-type human PKD2 in the *π*-conformation (PDB 5T4D, 3.0 Å^6^), the gain-of-function mutant F604P-PKD2 in the *α*-conformation (PDB 6D1W, 3.5 Å^8^), and PKD2L1 in the *α*-conformation (PDB 5Z1W, 3.4 Å^34^). Fourteen models were constructed (Table S1). For PKD2 systems, missing side chains and short loops were built and the required mutations were introduced using MODELLER 10.6,^44^ whereas the complete PKD2L1 structure was first obtained using Boltz-2 with PDB 5Z1W as a template.^45^ The resulting models covered residues 216–694 for the 5T4D-based systems, 216–682 for the 6D1W-based systems, and 101–563 for PKD2L1.

For the *α*-conformation of PKD2, an extended S6 helix was constructed using MODELLER by enforcing a standard *α*-helical conformation from residues 683–694, resulting in a model covering residues 216–694. To construct the alternative conformations, MODELLER was used following the approach described in a previous study of Na_V_ channels.^39^ Specifically, the initial PKD2 *α*-conformation model was constructed from PDB 5T4D by deleting F670 from both the template sequence and template structure. This caused the F670 residue in the resulting model to occupy the position of F671 in the *α*-conformation, thereby producing a standard *α*-helical register and shifting the downstream helical register by one residue. The PKD2L1 *π*-conformation model was constructed from the Boltz-2 *α*-conformation structure by introducing a gap in the template sequence immediately before F550. This caused F550 to occupy the position corresponding to the *π*-bulge and shifted the upstream helical register by one residue.

All models were then oriented in the membrane by alignment to the corresponding OPM entry^46^ and processed using PDB2PQR 3.7.1.^47^ Aspartate and glutamate residues were deprotonated, lysine and arginine residues were protonated, and all histidines in PKD2 were assigned the neutral HID tautomer. For PKD2L1, H258 was assigned the HIE tautomer and all other histidines were assigned HID. System construction was then performed using CHARMM-GUI.^48^ Chain termini were capped with acetyl and *N* -methylamide groups, and one disulfide bond was formed per chain (C331–C344 in PKD2 and C210–C223 in PKD2L1). Three *β*-D-GlcNAc residues were added to each subunit at N328, N362, and N375 in PKD2 and at the corresponding N177, N207, and N241 sites in PKD2L1. Each tetramer was embedded in a pure POPC bilayer using the CHARMM-GUI Membrane Builder,^49^ followed by solvation and neutralisation. All systems used a hexagonal simulation box with lateral dimensions of approximately 13.51 *×* 11.70 nm and a height of approximately 12.5–14.2 nm (Table S1). All systems contained 0.15 M KCl. For the ion-permeation simulations, additional systems were constructed with 0.15 M NaCl or 0.15 M CaCl_2_.

### Force fields and simulation parameters

Two sets of force fields were used in this study (Table S1). The first set comprised CHARMM36m for proteins,^50^ CHARMM36 parameters for lipids,^51^ CHARMM carbohydrate parameters,^52^ CHARMM-modified TIP3P water,^53^ and the standard CHARMM ion parameters.^54^ The electronic continuum correction (ECC) method was applied to this force-field set for more accurate channel conductance calculations. For monovalent ion conductance calculations, a charge-scaling factor of 0.78 was used based on a previous study of potassium channels,^32^ and the theoretical value of 0.75^31^ was additionally tested for K^+^. For Ca^2+^ conductance calculations, a charge-scaling factor of 0.87 was used based on a previous study of Ca_V_1 channels.^33^ The partial charges of all Asp, Glu, Lys, and Arg residues, as well as Cl*^−^* ions, were also scaled by the corresponding ion charge-scaling factor. The second force-field set comprised AMBER ff19SB for proteins,^55^ Lipid21 for lipids,^56^ and GLYCAM06j for glycans,^57^ together with OPC water^27^ and the 12-6-4 ion parameters.^58,59^ The force-field set used for each simulation is listed in Table S2.

All simulations were performed using GROMACS 2024.3^60^ patched with PLUMED 2.9.3,^61,62^ with a 2 fs integration time step (1 fs during the initial equilibration steps) and LINCS constraints applied to bonds involving hydrogen atoms.^63^ The Verlet neighbour list was updated every 100 steps. For CHARMM36m simulations, a 1.2 nm cutoff was used for both electrostatic and Lennard-Jones interactions. Lennard-Jones interactions were force-switched between 1.0 and 1.2 nm, and no long-range dispersion correction was applied. For AMBER simulations, a 0.9 nm cutoff was used for both electrostatic and Lennard-Jones interactions. Lennard-Jones interactions were truncated without switching, and long-range dispersion corrections were applied to the energy and pressure. Electrostatic interactions were calculated using the particle-mesh Ewald method,^64^ with a Fourier spacing of 0.12 nm and fourth-order interpolation.

The temperature was maintained at 310 K using the velocity-rescaling thermostat^65^ with a coupling time of 1 ps, applied separately to the solute, membrane, and solvent groups. The pressure was maintained at 1 bar using the C-rescale barostat, ^66^ with a coupling time of 5 ps and a compressibility of 4.5 *×* 10*^−^*^5^ bar*^−^*^1^, together with semi-isotropic pressure coupling. Centre-of-mass motion was removed every 1000 steps, separately for the solute–membrane and solvent groups.

### Simulation protocol

All systems were equilibrated using the standard CHARMM-GUI protocol, consisting of 5000 steps of energy minimisation followed by 250 ps of NVT and 1625 ps of NPT equilibration, with positional restraints released stepwise. The production simulations performed for each system are summarised in Table S2. In total, approximately 61 *µ*s of production simulations were performed. Unbiased simulations were performed without an external electric field. All pore-opening simulations and most restrained simulations were performed under a constant electric field of 0.02 V nm*^−^*^1^ (approximately 260 mV) applied along the membrane normal in the inward direction,^67^ with additional restrained simulations performed without the field as controls (Table S2).

In the restrained simulations, positional restraints with a force constant of 1000 kJ mol*^−^*^1^ nm*^−^*^2^ were applied either to all protein heavy atoms or to the protein backbone atoms (Table S2). The lower gate was opened with PLUMED by restraining C*α*–C*α* distances between the four equivalent gate residues (Fig. S4A,B). In the *α*-conformation, the two diagonal and four adjacent-subunit distances between the F676 or I680 residues were restrained, with each adjacent-subunit target set to the diagonal target divided by √2. The diagonal target was scanned from 14.5 to 19.0 Å for F676 and from 12.5 to 17.0 Å for I680 in 0.5 Å increments. With the original *α* model (up to residue 682), both scans were performed without and with additional lower-wall restraints on the diagonal (6.0 Å) and adjacent-subunit (4.2 Å) distances between the L641 carbonyl oxygen atoms of the selectivity filter, which prevented the filter from closing. With the extended model, the scans were performed without the wall restraints. For each target, the restraints were introduced over 10 ns by steered MD, with the force constants increased from zero to 3000 kJ mol*^−^*^1^ nm*^−^*^2^ for the diagonal distances and to 2000 kJ mol*^−^*^1^ nm*^−^*^2^ for the adjacent-subunit distances, followed by a 100 ns production simulation with the restraints held. In the *π*-conformation, the diagonal and adjacent-subunit L677 distances were restrained in the same way, with force constants of 3000 and 2000 kJ mol*^−^*^1^ nm*^−^*^2^, respectively, and the diagonal target was increased from 11.5 to 16.0 Å in 0.5 Å steps every 10 ns within a single 100 ns simulation. For the conductance simulations, the diagonal and adjacent-subunit L677 distances were brought to 14.5 and 10.3 Å, respectively, over 10 ns as described above and then held during the 500 ns production simulations with force constants of 3000 and 2000 kJ mol*^−^*^1^ nm*^−^*^2^.

In all pore-opening simulations, the C*α* atoms of residues 674–682 (674–680 in the original *α* model) were restrained along the membrane normal with a force constant of 1000 kJ mol*^−^*^1^ nm*^−^*^2^, and backbone dihedral restraints with a force constant of 200 kJ mol*^−^*^1^ rad*^−^*^2^ were applied to the same residues. A *χ*_1_ restraint with a force constant of 100 kJ mol*^−^*^1^ rad*^−^*^2^ was applied to the lower-gate residue F676 in the *α*-conformation and L677 in the *π*- conformation. For L677, the *χ*_1_ restraint used the angle observed in the 5MKE structure.^7^ An additional L677 distance scan was performed without the *χ*_1_ restraint on L677 (Fig. S16).

### Pore hydration analysis

The pore axis was defined in each frame as the line parallel to the membrane normal that passes through the *xy* centroid of the S6 C*α* atoms of all four subunits (residues 663–690 in PKD2 and the corresponding residues in PKD2L1). Water density was calculated for water oxygen atoms within 3 Å of the pore axis as a function of *z*, measured from the centroid of the L641 C*α* atoms (L521 in PKD2L1), with positive values towards the cytoplasm. A continuous water pathway was defined as an uninterrupted column of water molecules within 3 Å of the pore axis spanning the pore for at least 3 ns.

### Survey of TRP channel cryo-EM structures

All TRP channel EM entries in the PDB (retrieved August 2026, *n* = 684) were filtered to mammalian homotetramers with a modelled, fourfold-symmetric S6. The lower-gate residue was located in each chain by an S6 sequence motif, and each S6 was assigned as *π* or *α* according to which candidate residue faces the pore axis. For the 541 retained structures, C*α*– C*α* and closest heavy-atom distances between diagonally opposite subunits were averaged over the two pairs. Residues facing away from the pore were omitted. Residue assignments and exclusions for every entry are listed in Tables S3, S4 and S5.

### Ion permeation and conductance

Ion permeation was driven by a constant external electric field applied along the membrane normal.^67^ An electric field of 0.02 V nm*^−^*^1^ was used for monovalent ions, corresponding to approximately 260 mV across the simulation box according to *V* = *EL_z_*. Because no Ca^2+^ permeation events were observed at 0.02 V nm*^−^*^1^ in the 500ns AMBER simulations, a stronger field of 0.05 V nm*^−^*^1^ was used for Ca^2+^ to facilitate permeation, corresponding to approximately 650 mV, and the same field was used in the CHARMM simulations for consistency, corresponding to approximately 690 mV. The electric field was applied in both directions in separate simulations. For each ion and field direction, four independent replicas were performed, each consisting of 10 ns of gate opening followed by 500 ns of production.

Permeation events were counted using two planes defined in each frame by the L641 carbonyl oxygen atoms of the selectivity filter and by the I680 C*α* atoms below the lower gate. An inward event was counted when an ion entering from above the L641 plane subsequently crossed the I680 plane, and an outward event was counted when an ion entering from below the I680 plane reached the L641 plane. Pore excursions that did not complete the crossing, as well as apparent crossings caused by periodic boundary wrapping, were excluded.

The ionic current was calculated as

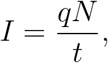

where *q* is the ionic charge, *N* is the net number of permeation events in the direction of the applied field, and *t* is the analysed simulation time, corresponding to the last 450 ns of each replica. The conductance was calculated as

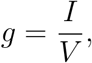

where *V* is the potential difference across the simulation box. Inward and outward conductances were obtained from simulations with the electric field applied in opposite directions. Conductances are reported as the mean *±* SD over four independent replicas.

### Software and packages

Trajectory analysis used MDAnalysis 2.8.0,^68^ NumPy 1.26.4, SciPy 1.15.2 and CHAP 0.9.1.^28^ Figures were prepared with Matplotlib 3.10.1, and molecular structures were visualised with Mol*^69^ and VMD 1.9.4a57.^70^

## Author contributions

C.Z. and J.M. designed research; C.Z. performed research; C.Z. analyzed data; W.K. and J.M. supervised research; C.Z. wrote the paper; and W.K. and J.M. reviewed and edited the paper.

## Data, Materials, and Software Availability

All simulation inputs and analysis code are available on GitHub at https://github.com/ michellab/PKD2_Function_In_Pi. The repository contains the starting structures, topologies and force-field parameter files, GROMACS .mdp files and PLUMED input files for all simulations listed in Table S2, the scripts used to analyse pore hydration, ion permeation and the TRP channel structure survey, and the representative structures shown in the figures. Owing to their size, the full simulation trajectories are not included, but trajectories supporting the main conclusions, with a reduced number of frames, are archived on Zenodo (DOI: 10.5281/zenodo.23946105).

## Acknowledgments

C.Z. thanks UKRI EPSRC (Project 2871271) for PhD scholarship funding. C.Z. and J.M. thank the HECBioSim Consortium (EPSRC grant EP/X035603/1) for providing access to the ARCHER2 UK National Supercomputing Service.

## Notes

https://github.com/michellab/PKD2_Function_In_Pi

https://dx.doi.org/10.5281/zenodo.23946105

### Competing Interest Statement

The authors have declared no competing interest.

