## Supporting Information for "TRPP Ion Channels Function in the *π*-Conformation Rather Than the *α*-Conformation"

### Supporting Information: TRPP Ion Channels Function in the $\pi$-Conformation Rather Than the $\alpha$-Conformation

### Supplementary figures

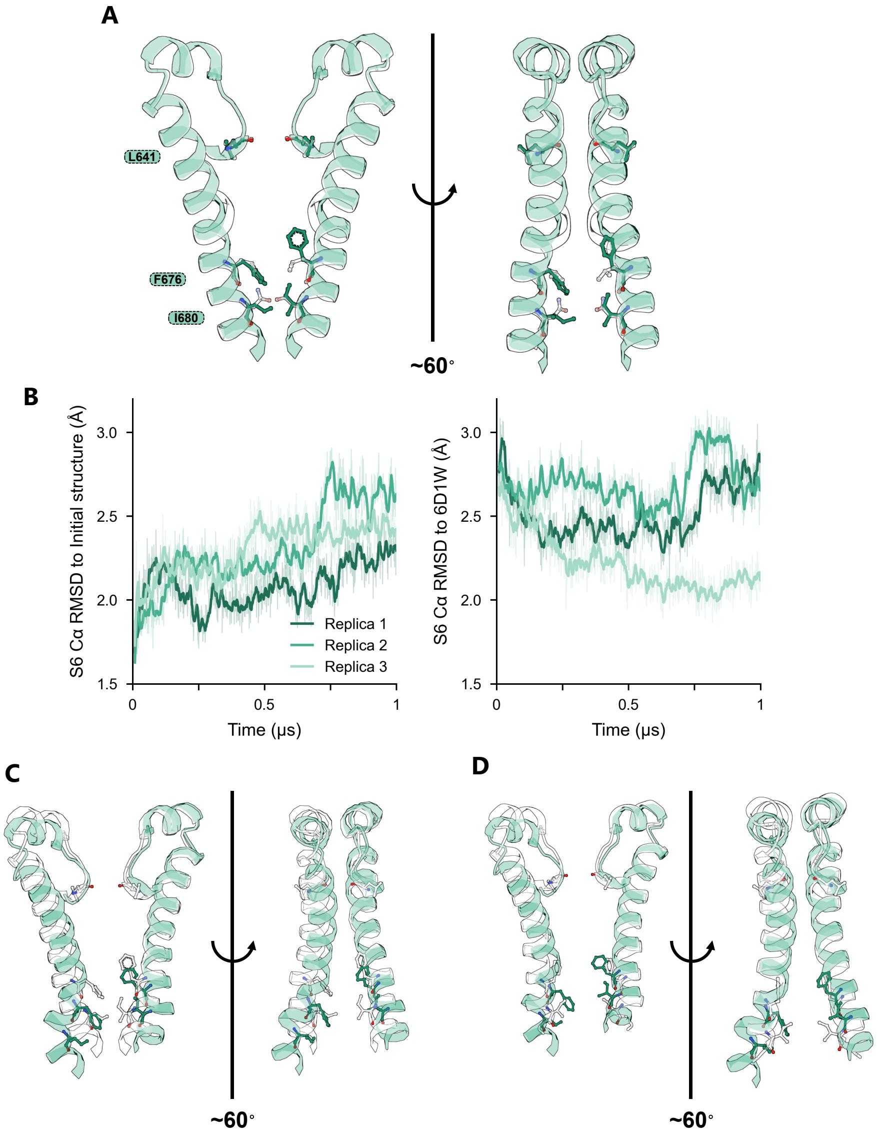


Figure S1: **Relaxation of the WT-$\alpha^{\mathrm{HM}}$ homology model.** (**A**) WT-$\alpha^{\mathrm{HM}}$ model (residues 640–684) constructed by homology modelling with a gap edit in the template (see Methods), superposed on its template, the $\pi$-conformation cryo-EM structure of WT-PKD2 (PDB 5T4D). Two opposing subunits are shown for clarity, in two views rotated by approximately 60°. (**B**) C$\alpha$ RMSD of S6 relative to the initial model (left) and to PDB 6D1W (right) over three 1 $\mu$s replicas. Thin lines show the raw data and thick lines the 10-ns running mean. (**C**) Final snapshot of replica 3 superposed on the white initial model. (**D**) The same snapshot superposed on the $\alpha$-conformation cryo-EM structure of F604P-PKD2 (PDB 6D1W).

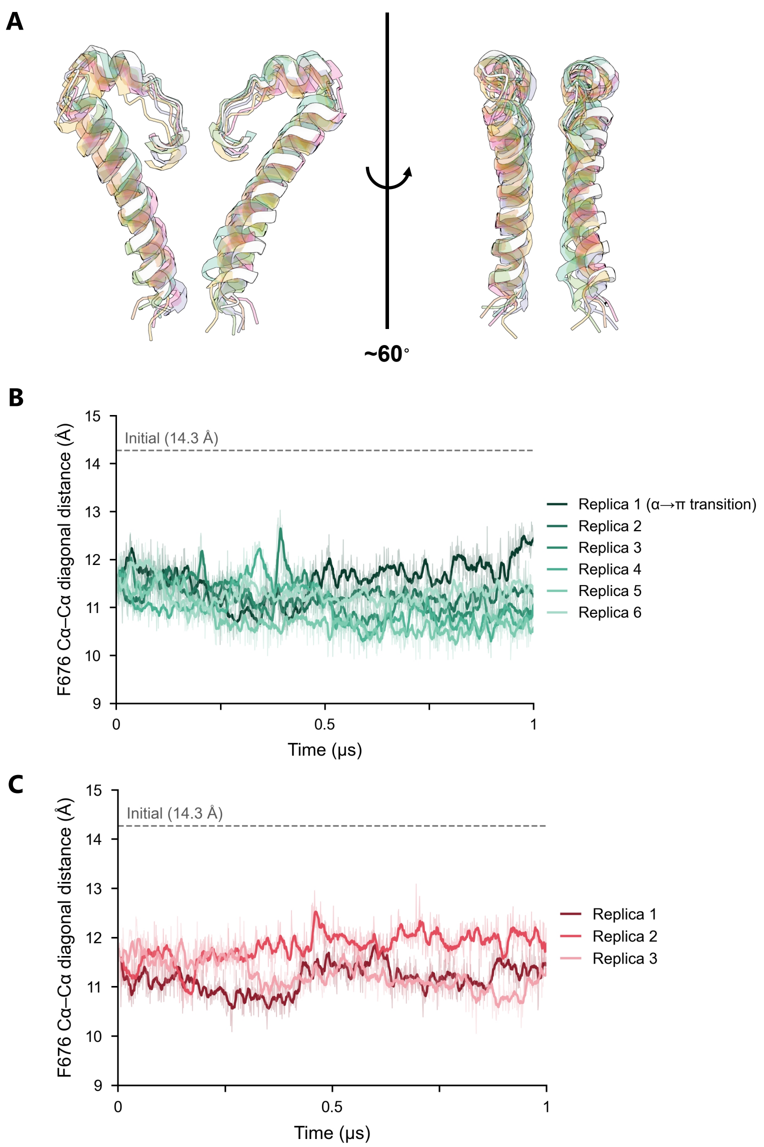


Figure S2: **Unbiased simulations of the WT-$\alpha$ and F604P-$\alpha$ models.** (**A**) Final snapshot of six 1 $\mu$s replicas superposed on the white initial WT-$\alpha$ model. (**B**) Diagonal C$\alpha$–C$\alpha$ distance between the F676 residues of WT-$\alpha$. Replica 1 contains the spontaneous $\alpha$-to-$\pi$ transition shown in Fig. 2B. (**C**) Diagonal C$\alpha$–C$\alpha$ distance between the F676 residues of F604P-$\alpha$ over three 1 $\mu$s replicas. Thin lines show the raw data and thick lines the 10-ns running mean.

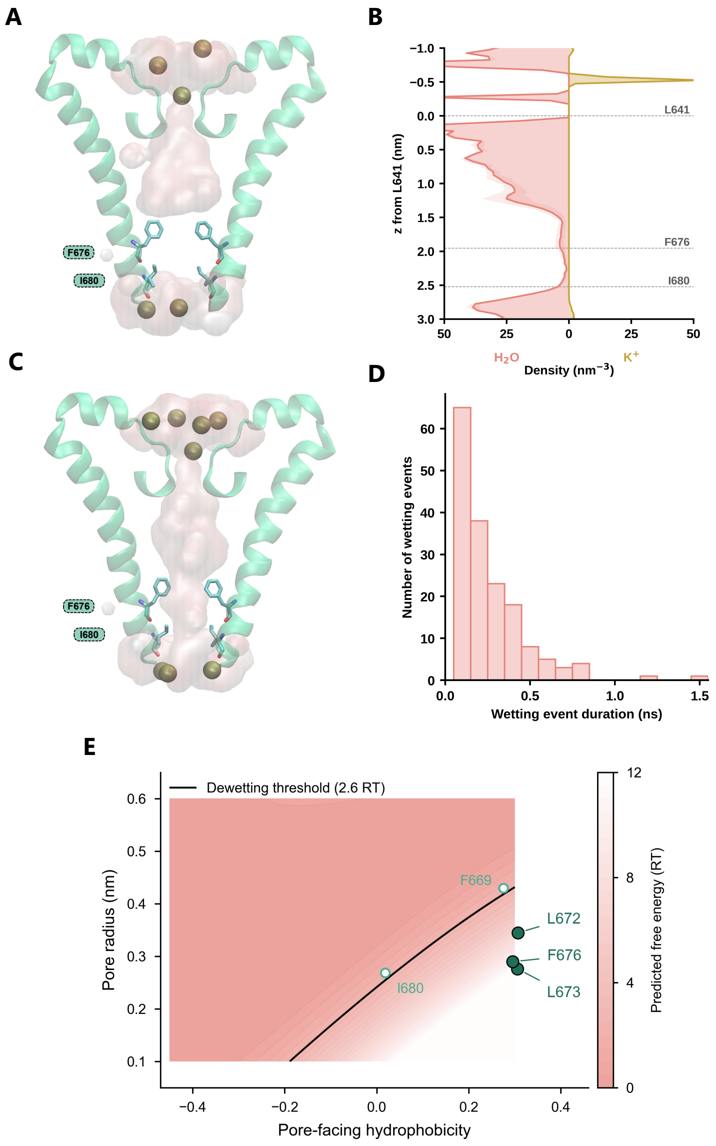


Figure S3: **Restrained simulations and CHAP prediction of the $\alpha$-conformation.** (**A,C**) Representative snapshot of WT-$\alpha$ with the protein heavy atoms restrained to the initial model with a dewetted (A) or hydrated (C) pore, simulated with CHARMM36m and TIP3P water under an applied electric field. Pore water is shown as a pink surface and $K^+$ ions as spheres. (**B**) Water (left) and $K^+$ (right) density within 3 Å of the pore axis, averaged over four 100-ns replicas (shading, SD across replicas). $z$ is measured from the selectivity filter (L641 C$\alpha$), with positive values towards the cytoplasm. (**D**) Duration of lower-gate wetting events. A frame is counted as wet when a continuous water column spans the gate from F676 to I680, with every 3 Å slice along the pore axis holding at least one water molecule within 3 Å of the axis. In total, 166 events were observed across four replicas (frames saved every 0.1 ns). (**E**) CHAP prediction for the initial $\alpha$-conformation structure. Pore radius is plotted against the pore-facing hydrophobicity of each pore-lining residue, with the background showing the predicted free energy of hydration. Filled circles mark residues predicted to be dewetted.

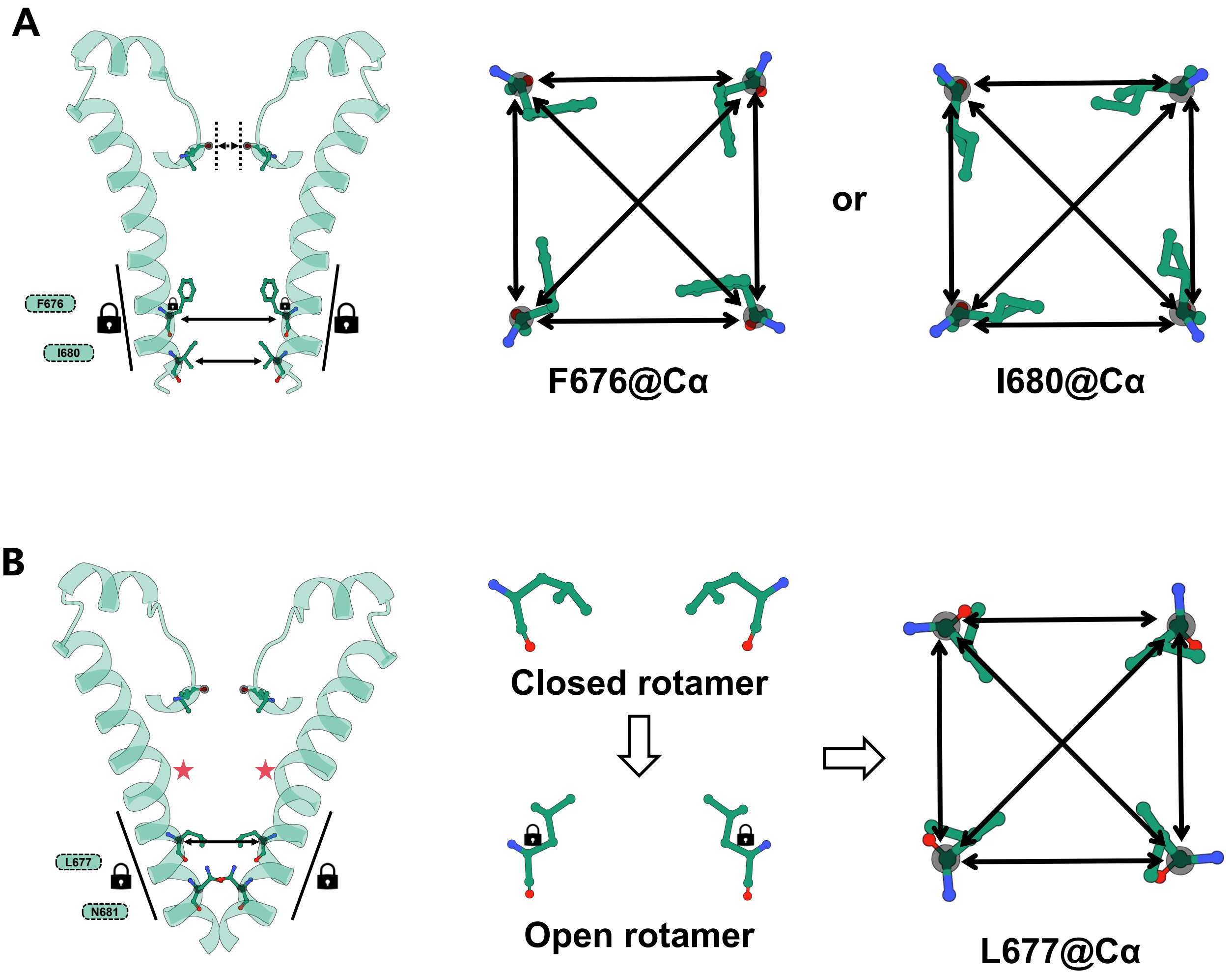


Figure S4: **Restraints used to open the lower gate.** (**A**) Restraints used to open the $\alpha$-conformation. Left, side view showing the C$\alpha$–C$\alpha$ distance restraints at F676 or I680 (arrows), the C$\alpha$ $z$-coordinate and backbone dihedral restraints on residues 674–680 (674–682 in the extended model, large locks) and the $\chi_1$ restraint on F676 (small locks). The dotted lines at the selectivity filter mark the lower-wall restraints on the L641 O distances, which were applied only in the scans of the original $\alpha$ model. Right, top view showing the two diagonal and four adjacent-subunit C$\alpha$–C$\alpha$ distances between the four F676 or I680 residues, which were steered to and then held at their target values. (**B**) Restraints used to open the $\pi$-conformation. Left, side view showing the C$\alpha$–C$\alpha$ distance restraints at L677 (arrows), the C$\alpha$ $z$-coordinate and backbone dihedral restraints on residues 674–682 (large locks) and the $\pi$-bulges (red stars). Middle, the closed rotamer of L677 (PDB 5T4D) and the open-like rotamer (PDB 5MKE) to which the $\chi_1$ angle was restrained (small locks). Right, top view showing the two diagonal and four adjacent-subunit C$\alpha$–C$\alpha$ distances between the four L677 residues, which were steered to and then held at their target values.

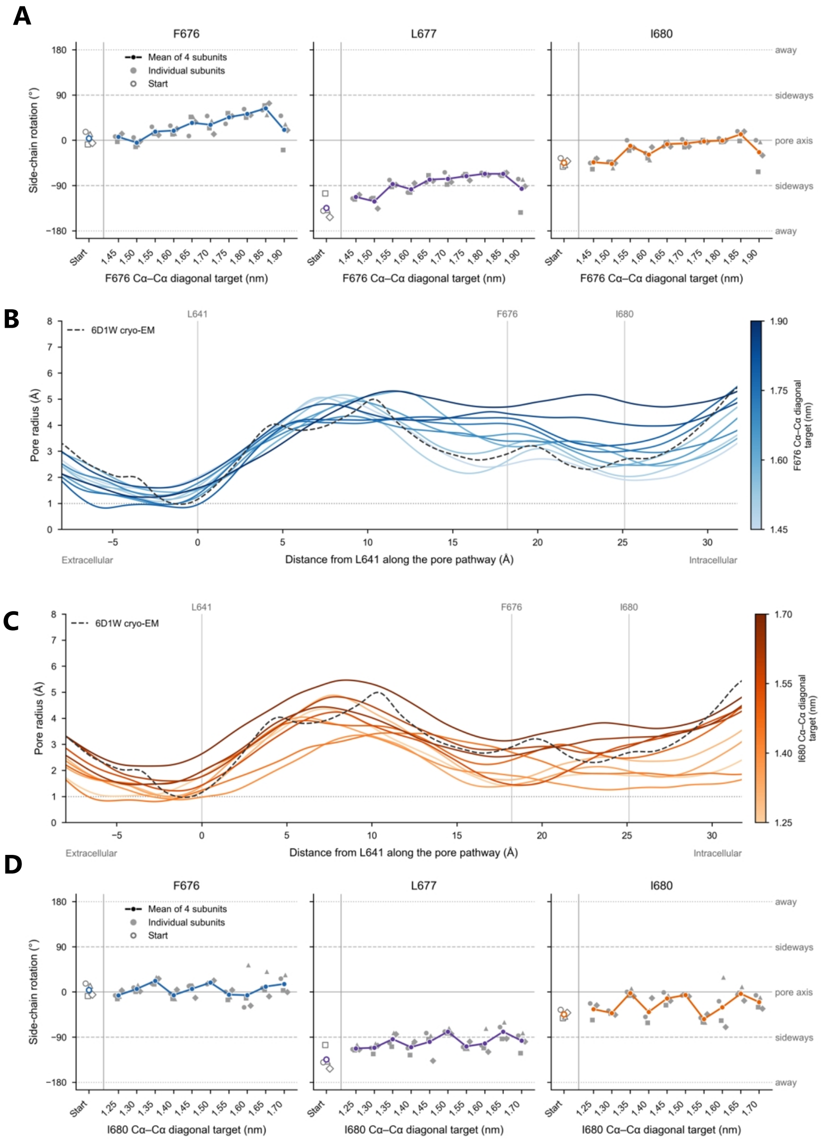


Figure S5: **F676 and I680 distance scans of the original $\alpha$-conformation model.** (**A**) Side-chain rotation of F676, L677 and I680 as a function of the diagonal F676 C$\alpha$–C$\alpha$ target distance. 0° indicates a side chain pointing towards the pore axis, ±90° sideways and ±180° away from the pore. The pore axis was defined in each frame by the $xy$ centroid of the F600, F676 and I680 C$\alpha$ atoms. Each subunit was averaged over the last 10 ns of each target distance. (**B**) Pore radius along the pore for each target of the F676 scan, averaged over the last 10 ns, with distance measured from L641 C$\alpha$. (**C**) As in B for the I680 scan. (**D**) As in A for the I680 scan.

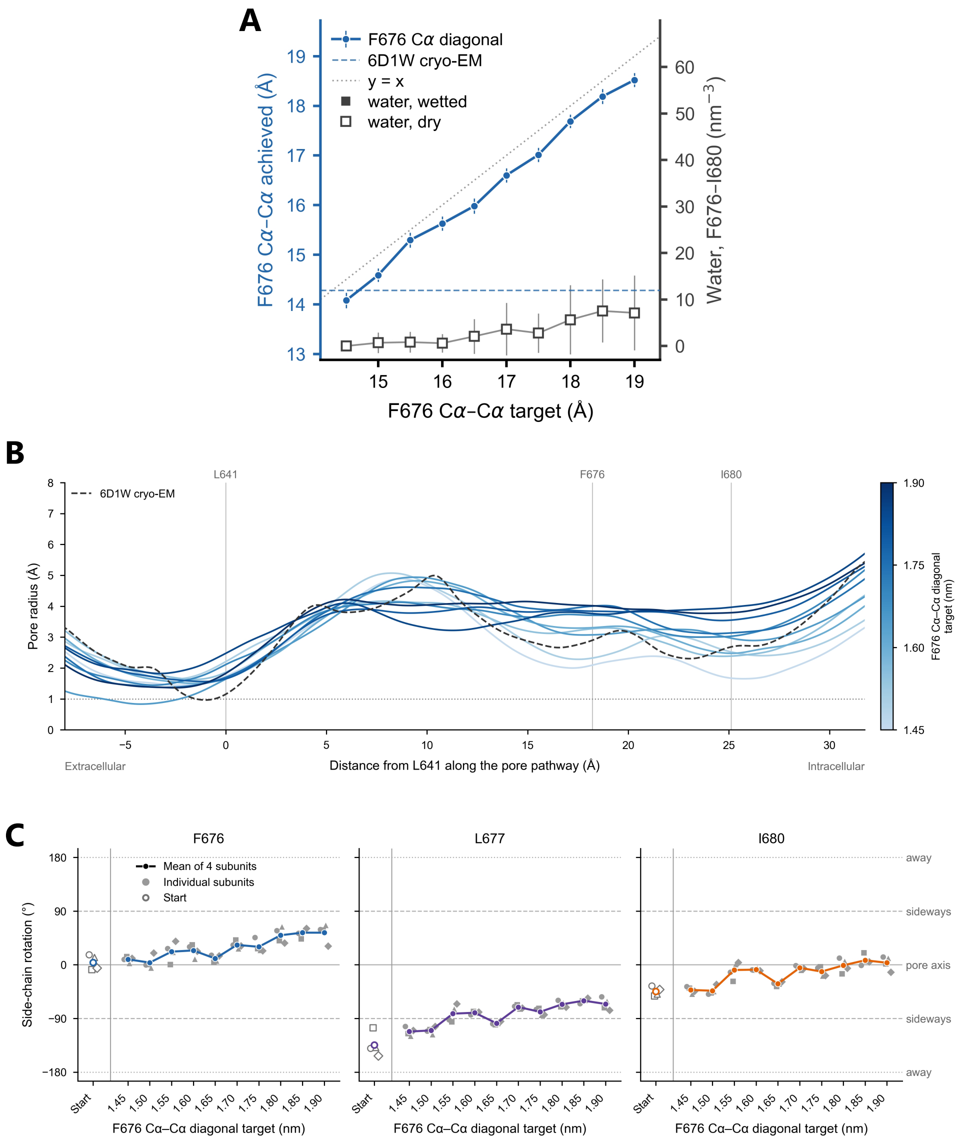


Figure S6: **F676 distance scan of the original $\alpha$-conformation model with a lower-wall restraint on the selectivity filter.** (**A**) Achieved diagonal F676 C$\alpha$–C$\alpha$ distance (left axis) and water density between the F676 and I680 C$\alpha$ planes (right axis) as a function of the target distance. Mean ± SD values were calculated over the last 10 ns of each target. The dashed line marks the value in the cryo-EM structure (PDB 6D1W, 14.28 Å) and the dotted line indicates $y = x$. Filled and open squares mark hydrated and dry pores. No continuous water pathway formed at any target distance. (**B**) Pore radius along the pore for each target of the F676 scan, averaged over the last 10 ns, with distance measured from L641 C$\alpha$. (**C**) Side-chain rotation of F676, L677 and I680, defined as in Fig. S5A.

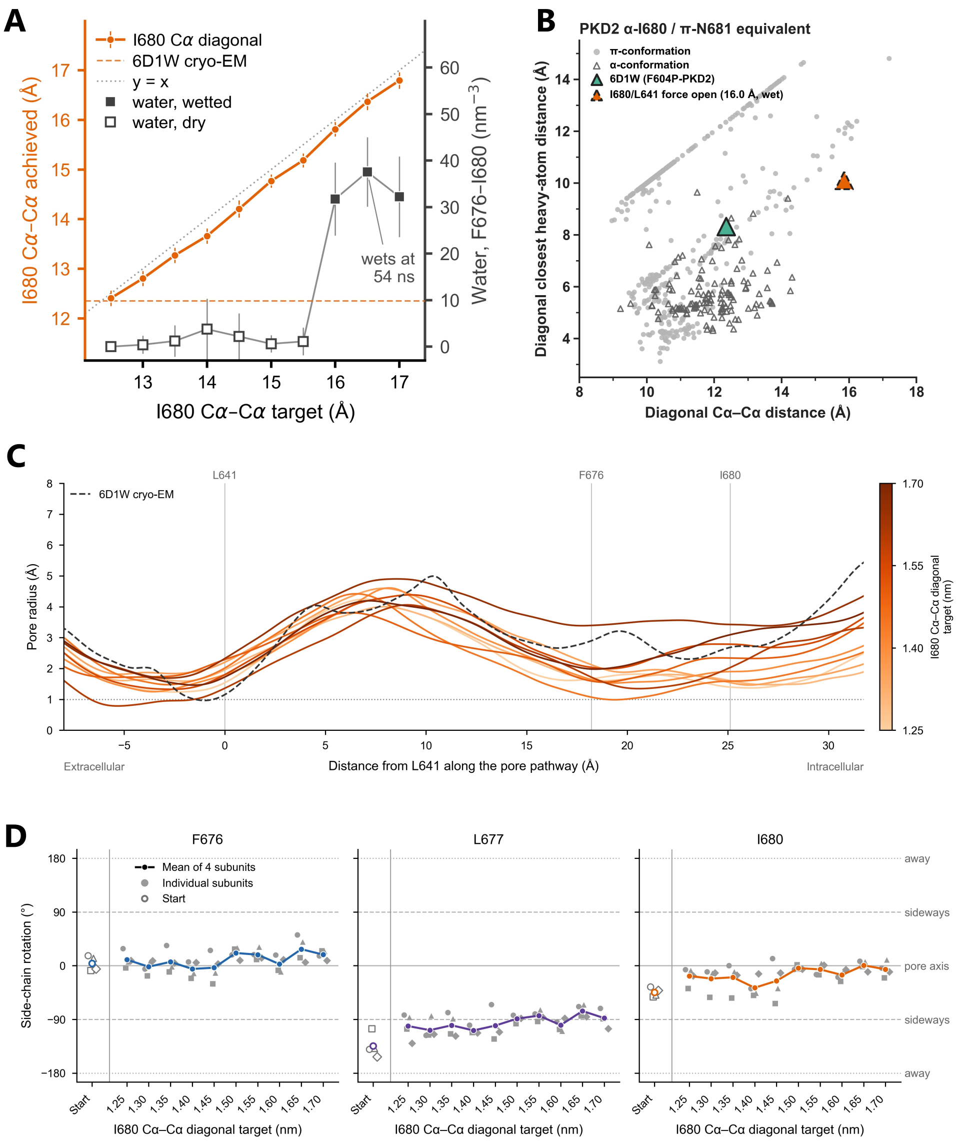


Figure S7: **I680 distance scan of the original $\alpha$-conformation model with a lower-wall restraint on the selectivity filter.** (**A**) Achieved diagonal I680 C$\alpha$–C$\alpha$ distance (left axis) and water density between the F676 and I680 C$\alpha$ planes (right axis) as a function of the target distance. Mean $\pm$ SD values were calculated over the last 10 ns of each target. The dashed line marks the value in the cryo-EM structure (PDB 6D1W, 12.35 Å) and the dotted line indicates $y = x$. Filled and open squares mark hydrated and dry pores. The pore first became hydrated at a target distance of 16.0 Å. (**B**) Diagonal closest heavy-atom distance versus diagonal C$\alpha$–C$\alpha$ distance between the $\alpha$-I680/$\pi$-N681-equivalent residues in mammalian TRP cryo-EM structures, with PDB 6D1W highlighted. The values for the hydrated $\alpha$-conformation were calculated from the final frame of the first hydrated simulation. (**C**) Pore radius along the pore for each target of the I680 scan, averaged over the last 10 ns, with distance measured from L641 C$\alpha$. (**D**) Side-chain rotation of F676, L677 and I680, defined as in Fig. S5A.

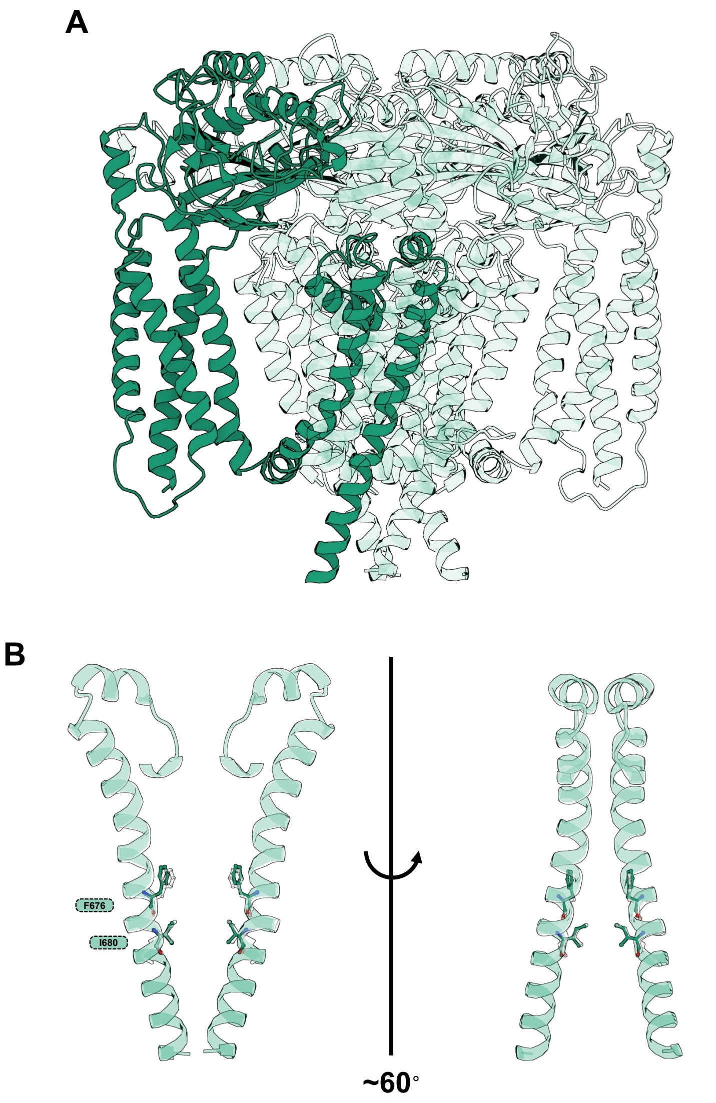


Figure S8: **Extended $\alpha$-conformation model.** (**A**) Overall structure of the WT-$\alpha$ model extended to residue 694, with one subunit highlighted. (**B**) Pore-lining S6 helices of two opposing subunits in two views rotated by approximately 60°, with F676 and I680 shown as sticks, superposed on the white original model (up to residue 682).

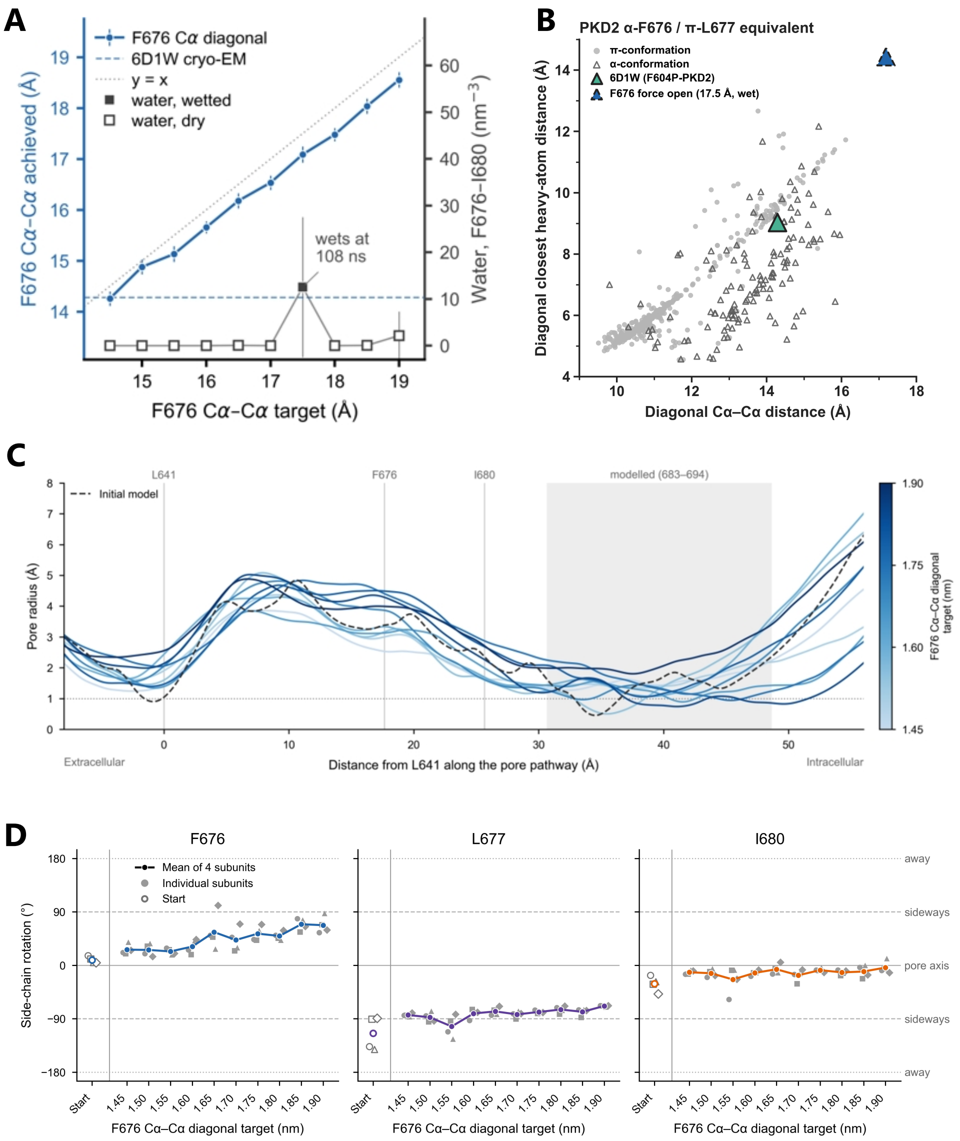


Figure S9: **F676 distance scan of the extended $\alpha$-conformation model.** (**A**) Achieved diagonal F676 C$\alpha$–C$\alpha$ distance (left axis) and water density between the F676 and I680 C$\alpha$ planes (right axis) as a function of the target distance. Mean ± SD values were calculated over the last 10 ns of each target. The dashed line marks the value in the cryo-EM structure (PDB 6D1W, 14.28 Å) and the dotted line indicates $y = x$. Filled and open squares mark hydrated and dry pores. The pore first became hydrated at a target distance of 17.5 Å. (**B**) Diagonal closest heavy-atom distance versus diagonal C$\alpha$–C$\alpha$ distance between the $\alpha$-F676/$\pi$-L677-equivalent residues in mammalian TRP cryo-EM structures, with PDB 6D1W highlighted. The values for the hydrated $\alpha$-conformation were calculated from the final frame of the first hydrated simulation. (**C**) Pore radius along the pore for each target of the F676 scan, averaged over the last 10 ns, with distance measured from L641 C$\alpha$. The grey band marks residues 683–694, which are not resolved in the cryo-EM structure and were modelled. (**D**) Side-chain rotation of F676, L677 and I680, defined as in Fig. S5A.

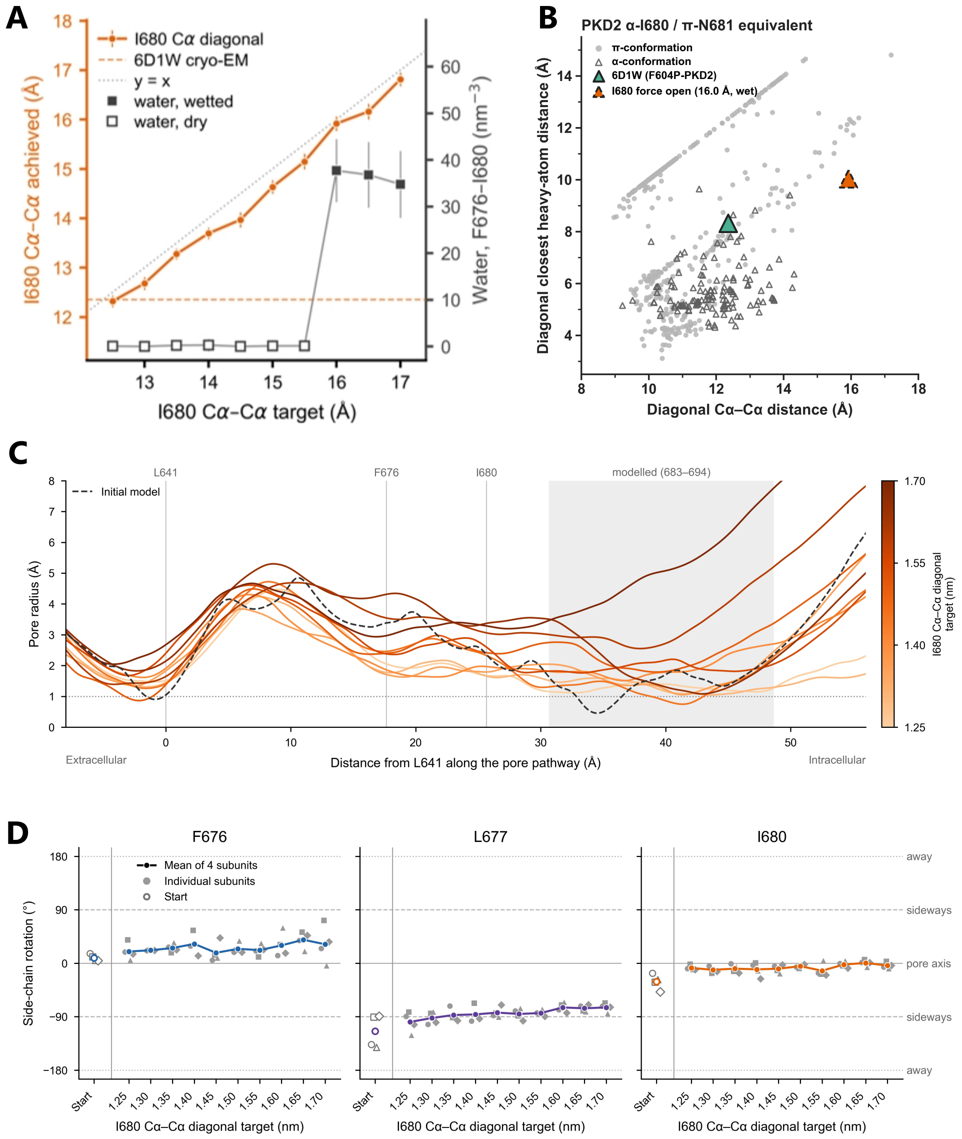


Figure S10: **I680 distance scan of the extended $\alpha$-conformation model.** (**A**) Achieved diagonal I680 C$\alpha$–C$\alpha$ distance (left axis) and water density between the F676 and I680 C$\alpha$ planes (right axis) as a function of the target distance. Mean ± SD values were calculated over the last 10 ns of each target. The dashed line marks the value in the cryo-EM structure (PDB 6D1W, 12.35 Å) and the dotted line indicates $y = x$. Filled and open squares mark hydrated and dry pores. The pore first became hydrated at a target distance of 16.0 Å. (**B**) Diagonal closest heavy-atom distance versus diagonal C$\alpha$–C$\alpha$ distance between the $\alpha$-I680/$\pi$-N681-equivalent residues in mammalian TRP cryo-EM structures, with PDB 6D1W highlighted. The values for the hydrated $\alpha$-conformation were calculated from the final frame of the first hydrated simulation. (**C**) Pore radius along the pore for each target of the I680 scan, averaged over the last 10 ns, with distance measured from L641 C$\alpha$. The grey band marks residues 683–694, which are not resolved in the cryo-EM structure and were modelled. (**D**) Side-chain rotation of F676, L677 and I680, defined as in Fig. S5A.

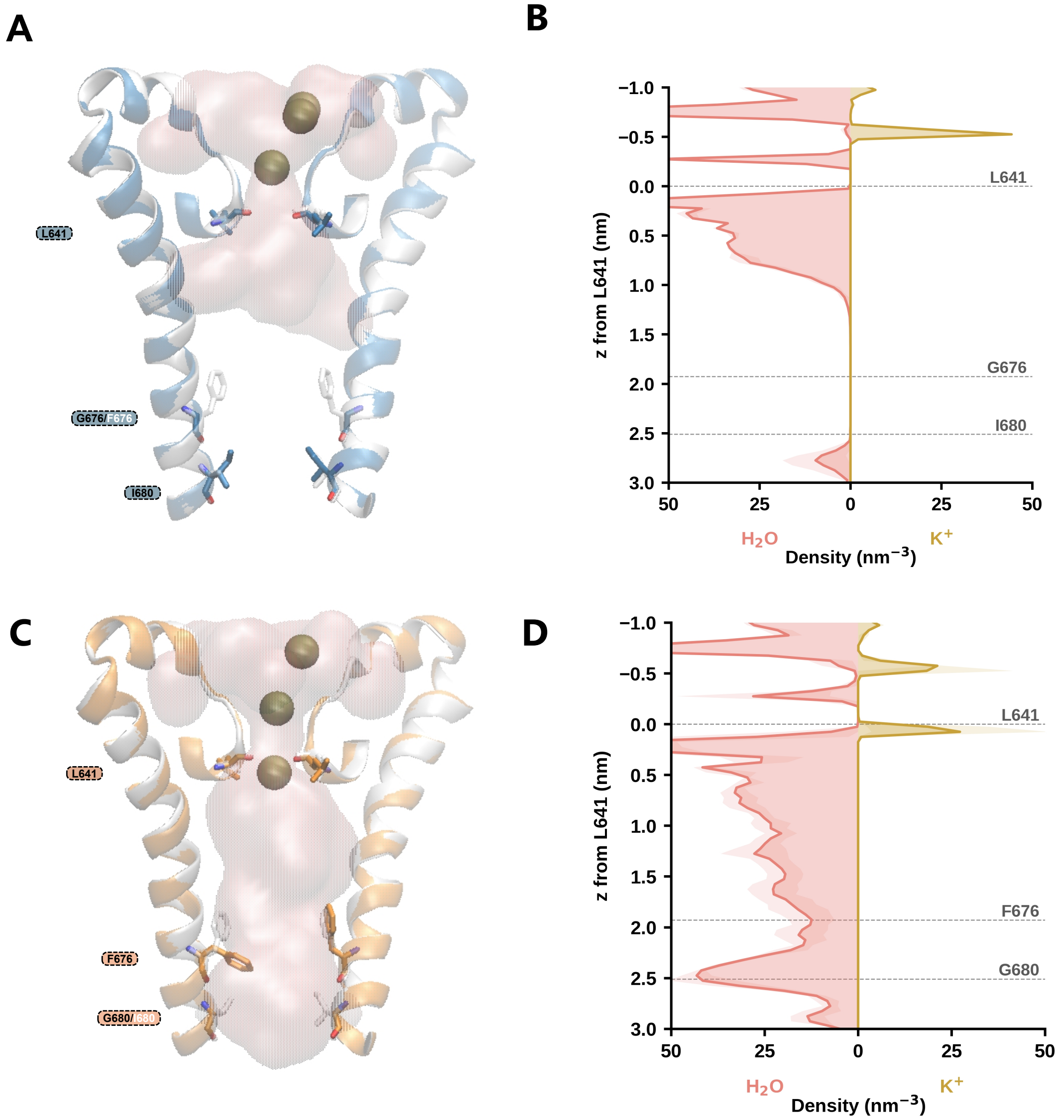


Figure S11: **F676G and I680G mutants of the extended $\alpha$-conformation model.** (**A,C**) Representative snapshots of F676G (A) and I680G (C) with the protein backbone atoms restrained to the initial model under an applied electric field, superposed on the white initial model. Pore water is shown as a pink surface and $K^+$ ions as spheres. (**B,D**) Water (left) and $K^+$ (right) density within 3 Å of the pore axis for F676G (B) and I680G (D), averaged over three 100-ns replicas (shading, SD across replicas). $z$ is measured from the selectivity filter (L641 C$\alpha$), with positive values towards the cytoplasm. As in the original model, I680G but not F676G results in pore hydration.

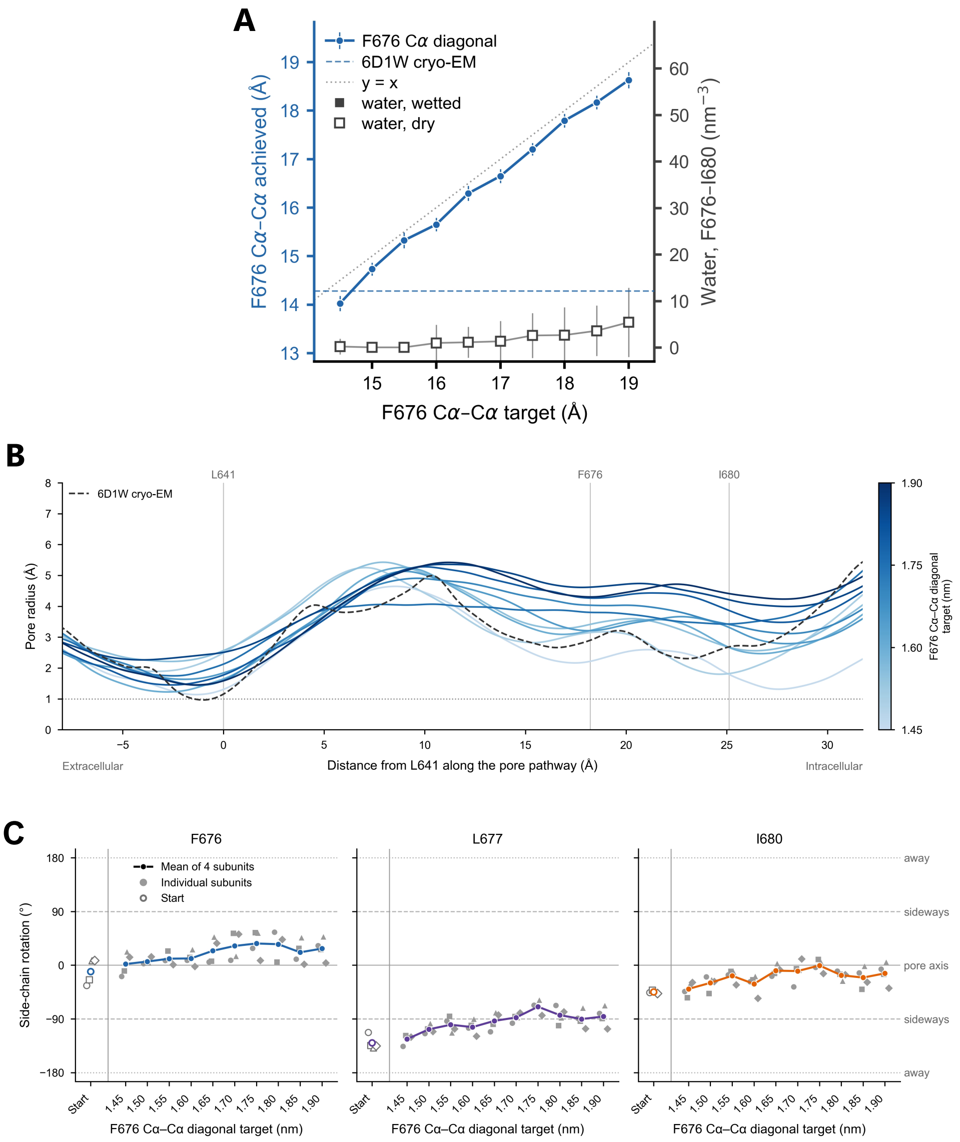


Figure S12: **F676 distance scan of the original $\alpha$-conformation model with TIP3P water.** (**A**) Achieved diagonal F676 C$\alpha$–C$\alpha$ distance (left axis) and water density between the F676 and I680 C$\alpha$ planes (right axis) as a function of the target distance. Mean ± SD values were calculated over the last 10 ns of each target. The dashed line marks the value in the cryo-EM structure (PDB 6D1W, 14.28 Å) and the dotted line indicates $y = x$. Filled and open squares mark hydrated and dry pores. No continuous water pathway formed at any target distance. (**B**) Pore radius along the pore for each target of the F676 scan, averaged over the last 10 ns, with distance measured from L641 C$\alpha$. (**C**) Side-chain rotation of F676, L677 and I680, defined as in Fig. S5A.

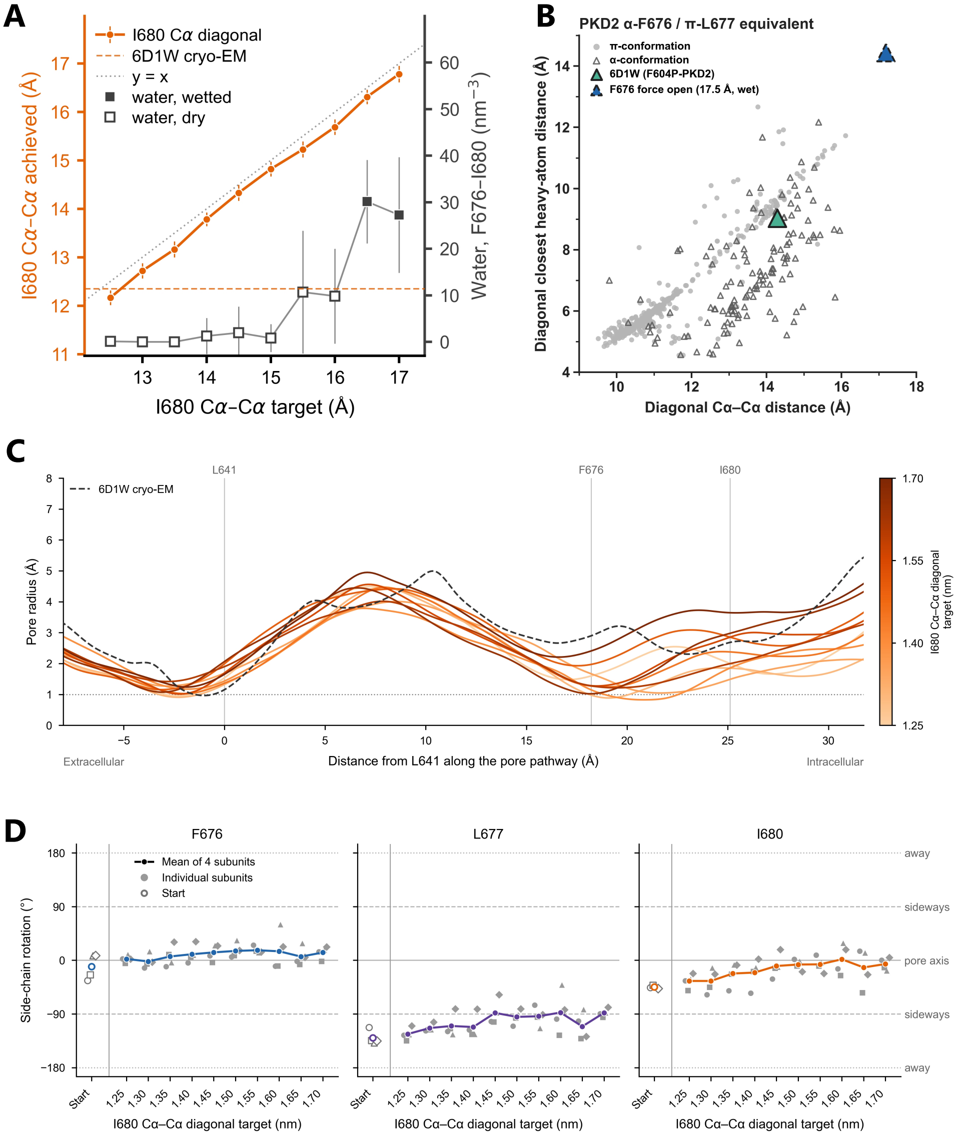


Figure S13: **I680 distance scan of the original $\alpha$-conformation model with TIP3P water.** (**A**) Achieved diagonal I680 C$\alpha$–C$\alpha$ distance (left axis) and water density between the F676 and I680 C$\alpha$ planes (right axis) as a function of the target distance. Mean ± SD values were calculated over the last 10 ns of each target. The dashed line marks the value in the cryo-EM structure (PDB 6D1W, 12.35 Å) and the dotted line indicates $y = x$. Filled and open squares mark hydrated and dry pores. The pore first became hydrated at a target distance of 16.5 Å. (**B**) Diagonal closest heavy-atom distance versus diagonal C$\alpha$–C$\alpha$ distance between the $\alpha$-I680/$\pi$-N681-equivalent residues in mammalian TRP cryo-EM structures, with PDB 6D1W highlighted. The values for the hydrated $\alpha$-conformation were calculated from the final frame of the first hydrated simulation. (**C**) Pore radius along the pore for each target of the I680 scan, averaged over the last 10 ns, with distance measured from L641 C$\alpha$. (**D**) Side-chain rotation of F676, L677 and I680, defined as in Fig. S5A.

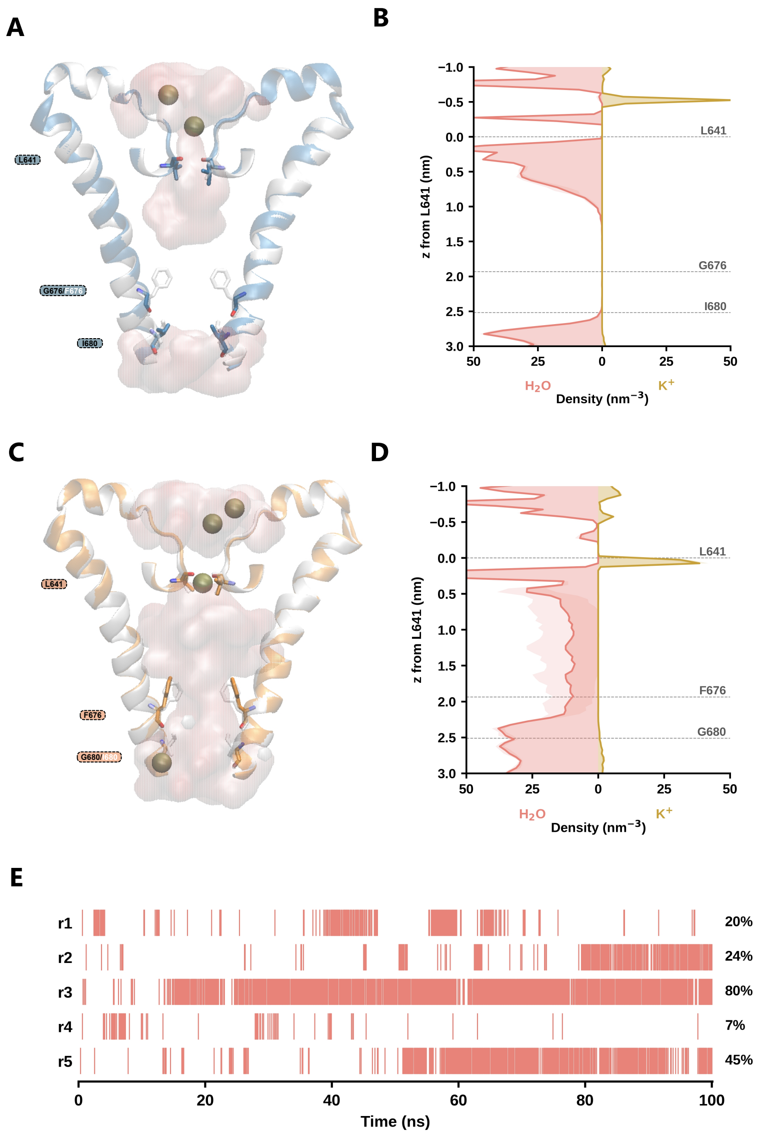


Figure S14: **F676G and I680G mutants of the original $\alpha$-conformation model with TIP3P water.** (**A**) Representative snapshot of F676G with the protein backbone atoms restrained to the initial model under an applied electric field, superposed on the white initial model. (**B**) Water (left) and $K^+$ (right) density within 3 Å of the pore axis for F676G, averaged over three 100-ns replicas (shading, SD across replicas). (**C**) Representative snapshot of I680G under the same conditions. (**D**) As in B for I680G, averaged over five 100-ns replicas. (**E**) Lower-gate wetting events in each replica of I680G. The percentage of time with a continuous water column across the lower gate is given for each replica. Pore water is shown as a pink surface and $K^+$ ions as spheres. $z$ is measured from the selectivity filter (L641 C$\alpha$), with positive values towards the cytoplasm.

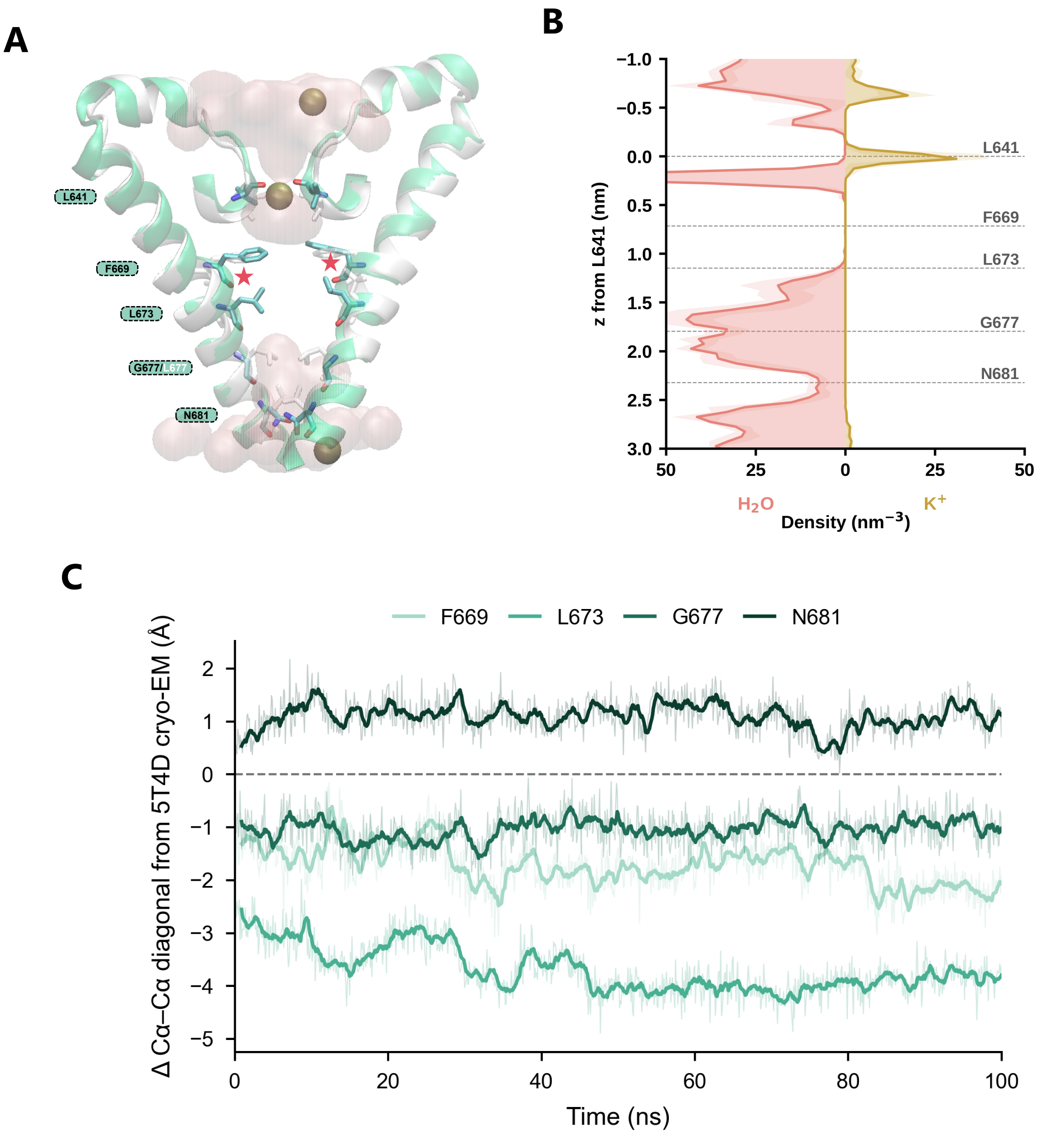


Figure S15: **Unbiased simulation of L677G-$\pi$.** (**A**) Representative snapshot of L677G-$\pi$ during unbiased MD without an applied electric field, superposed on the white initial model (Fig. 4A). $\pi$-bulges are marked by red stars. Pore water is shown as a pink surface and $K^+$ ions as spheres. (**B**) Water (left) and $K^+$ (right) density within 3 Å of the pore axis, averaged over the 100-ns simulation (shading, SD over five 20-ns blocks). $z$ is measured from the selectivity filter (L641 C$\alpha$), with positive values towards the cytoplasm. The region between G677 and N681 is hydrated, whereas the pore between F669 and L673 is dry. (**C**) Change in the mean of the two diagonal C$\alpha$–C$\alpha$ distances at F669, L673, G677 and N681 relative to the cryo-EM structure (PDB 5T4D). The dashed line marks zero change. Thin lines show the raw data and thick lines the 1-ns running mean.

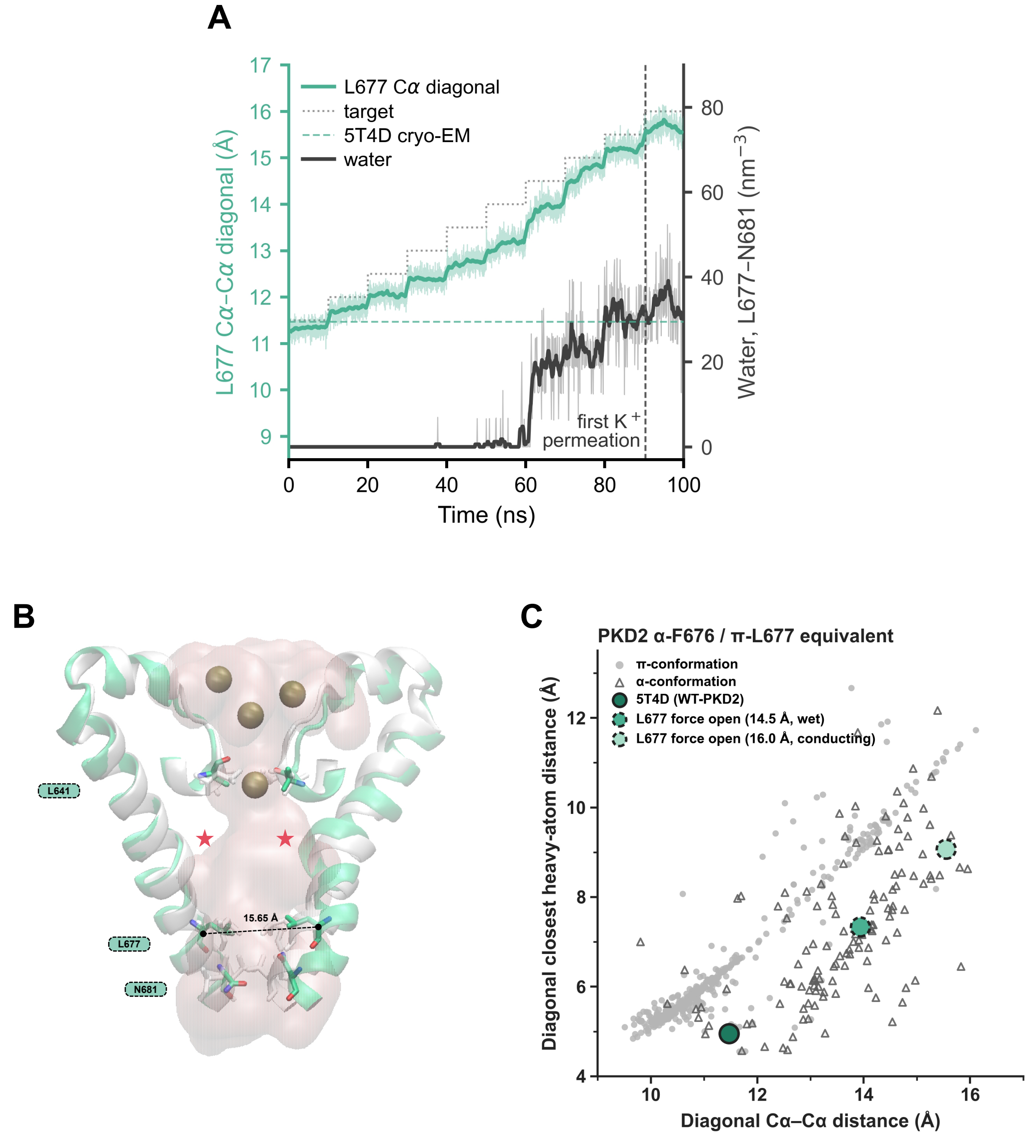


Figure S16: **Gate opening of the $\pi$-conformation without restraining L677 in the open-like rotamer.** (**A**) Stepwise increase of the diagonal L677 C$\alpha$–C$\alpha$ distance (left axis) and water density between the L677 and N681 C$\alpha$ planes (right axis). The dotted line shows the target distance and the horizontal dashed line the value in the cryo-EM structure (PDB 5T4D). The vertical dashed line marks the first $K^+$ permeation event. Thin lines show the raw data and thick lines the 1-ns running mean. (**B**) Representative snapshot at the first $K^+$ permeation event. $\pi$-bulges are marked by red stars. Pore water is shown as a pink surface and $K^+$ ions as spheres. (**C**) Diagonal closest heavy-atom distance versus diagonal C$\alpha$–C$\alpha$ distance between the L677-equivalent residues in mammalian TRP cryo-EM structures, with PDB 5T4D highlighted. The values at the first hydration (14.5 Å) and the first $K^+$ permeation (16.0 Å) of the scan in A are highlighted.

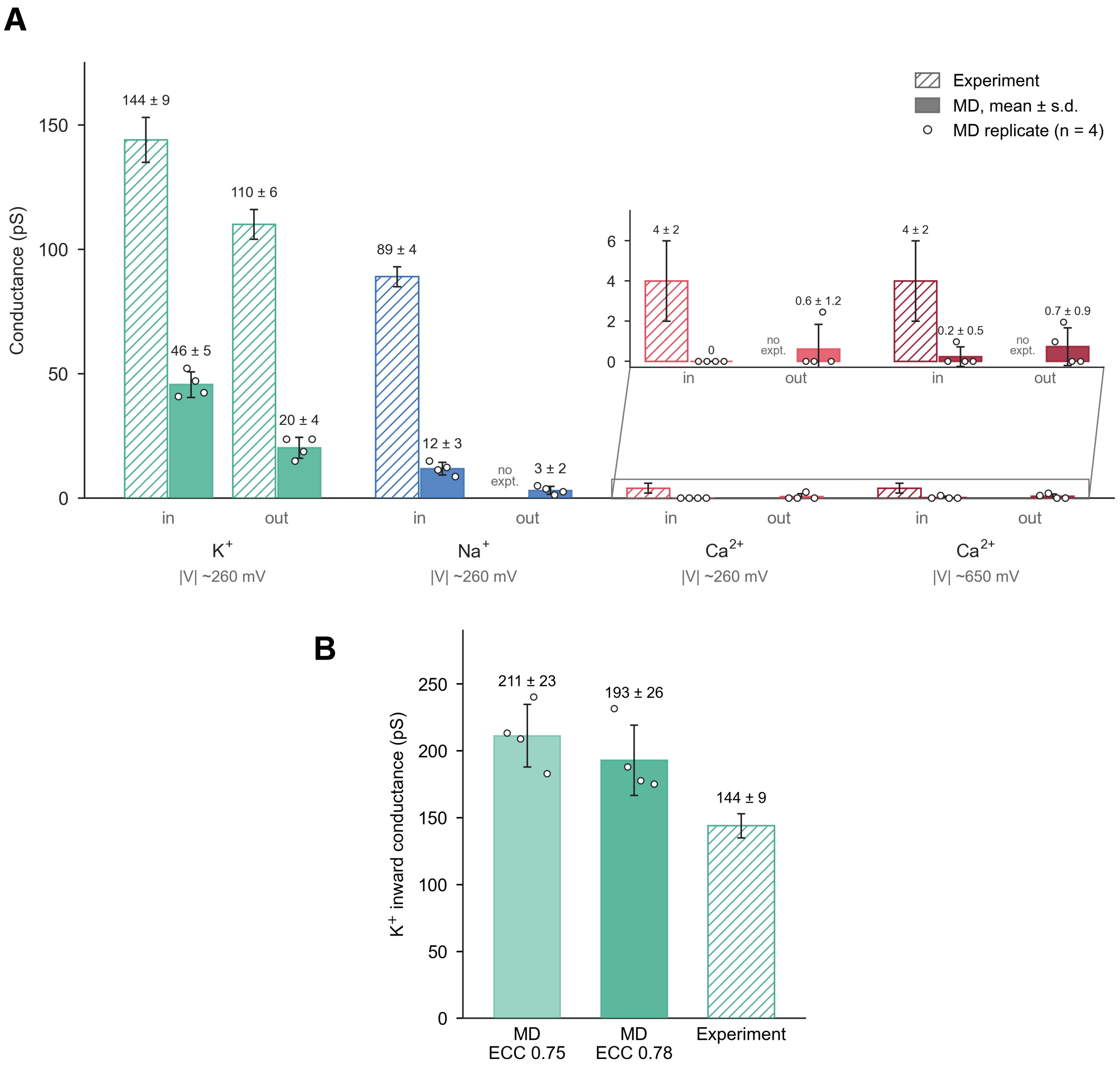


Figure S17: **Ion conductance of the opened $\pi$-conformation with different force fields and ECC scaling factors.** (**A**) Simulated conductance of $K^+$, $Na^+$ and $Ca^{2+}$ with AMBER ff19SB and OPC water (filled bars), compared with experimental values (hatched bars). Inward (in) and outward (out) conductances are shown at the indicated applied potentials $|V|$. Bars show the mean ± SD over four replicas and circles the individual replicas. The inset shows the $Ca^{2+}$ values on an expanded scale. (**B**) Inward $K^+$ conductance with CHARMM36m and ECC charge-scaling factors of 0.75 and 0.78, compared with experiment. Experimental values are as in Fig. 4G.

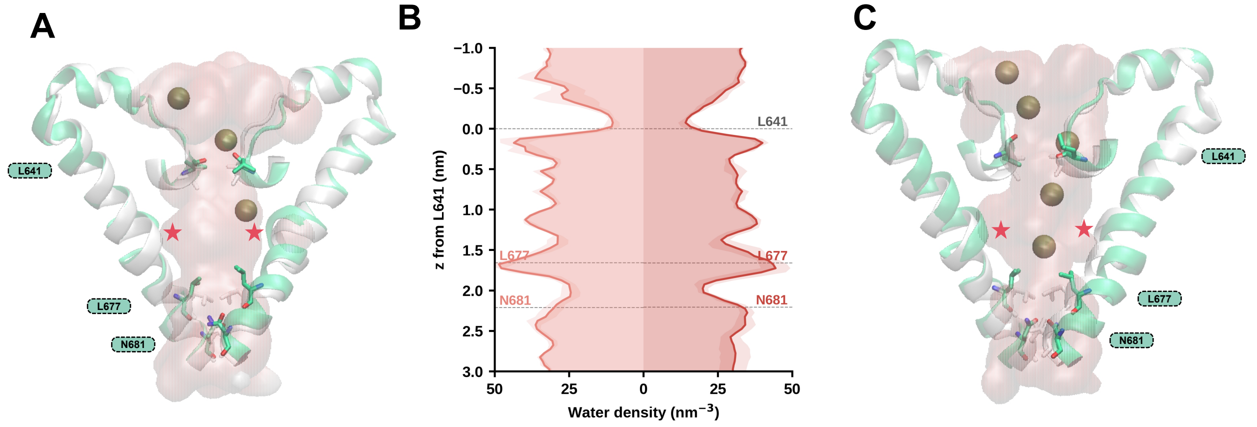


Figure S18: **Pore hydration of the opened $\pi$-conformation with OPC and TIP3P water.** (**A,C**) Representative snapshots of the $\pi$-conformation with the diagonal L677 C$\alpha$–C$\alpha$ distance restrained to 14.5 Å, simulated with OPC (A) and TIP3P (C) water. $\pi$-bulges are marked by red stars. Pore water is shown as a pink surface and $K^+$ ions as spheres. (**B**) Water density within 3 Å of the pore axis with OPC (left) and TIP3P (right) water, averaged over the last 450 ns of four replicas of the inward $K^+$ conductance simulations (shading, SD across replicas). $z$ is measured from the selectivity filter (L641 C$\alpha$), with positive values towards the cytoplasm. A continuous water pathway formed with both water models.

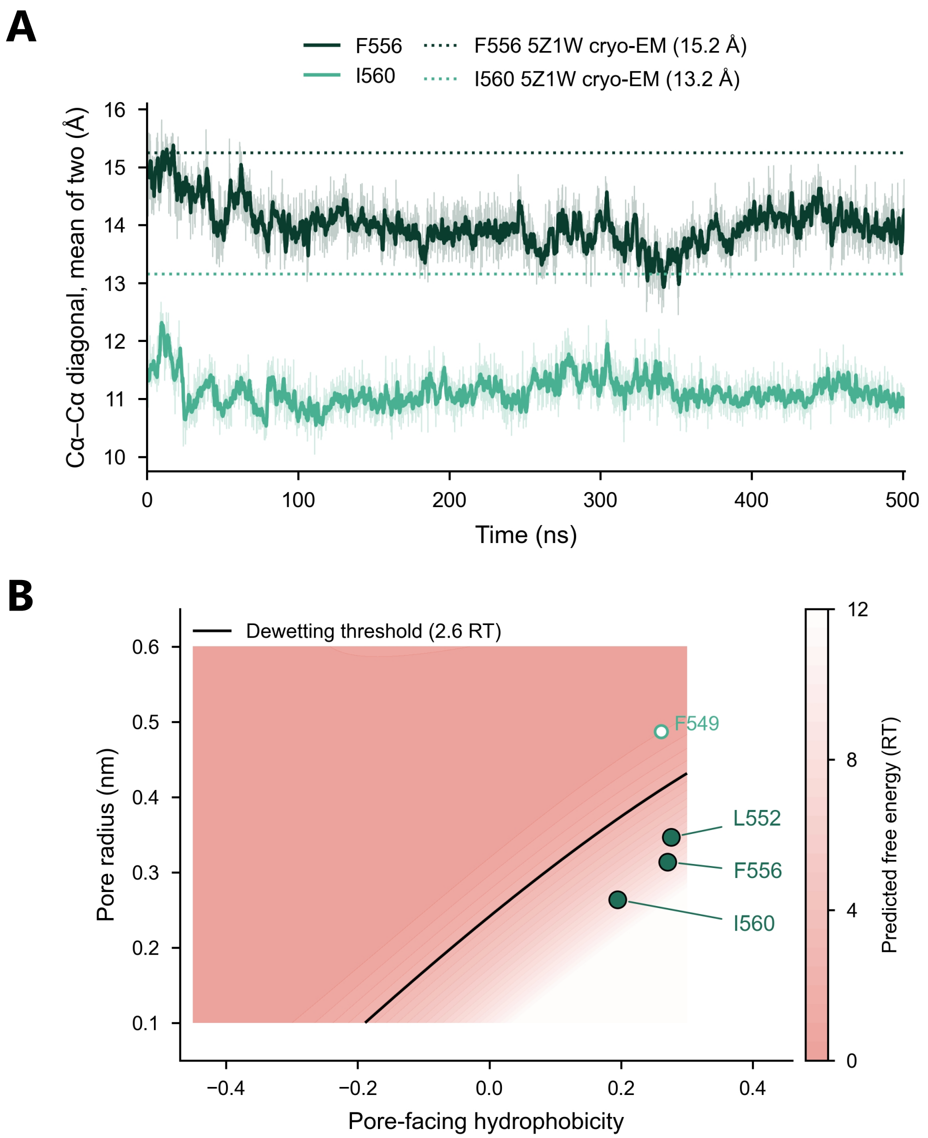


Figure S19: **Unbiased simulation and CHAP prediction of the PKD2L1 $\alpha$-conformation.** (**A**) Mean of the two diagonal C$\alpha$–C$\alpha$ distances at F556 and I560 during the 500 ns unbiased simulation of PKD2L1-$\alpha$ without an applied electric field. Dotted lines mark the values in the cryo-EM structure (PDB 5Z1W). Both distances decreased, indicating pore collapse. Thin lines show the raw data and thick lines the 1-ns running mean. (**B**) CHAP prediction for the initial PKD2L1-$\alpha$ model, shown as in Fig. S3E. Filled circles mark residues predicted to be dewetted.

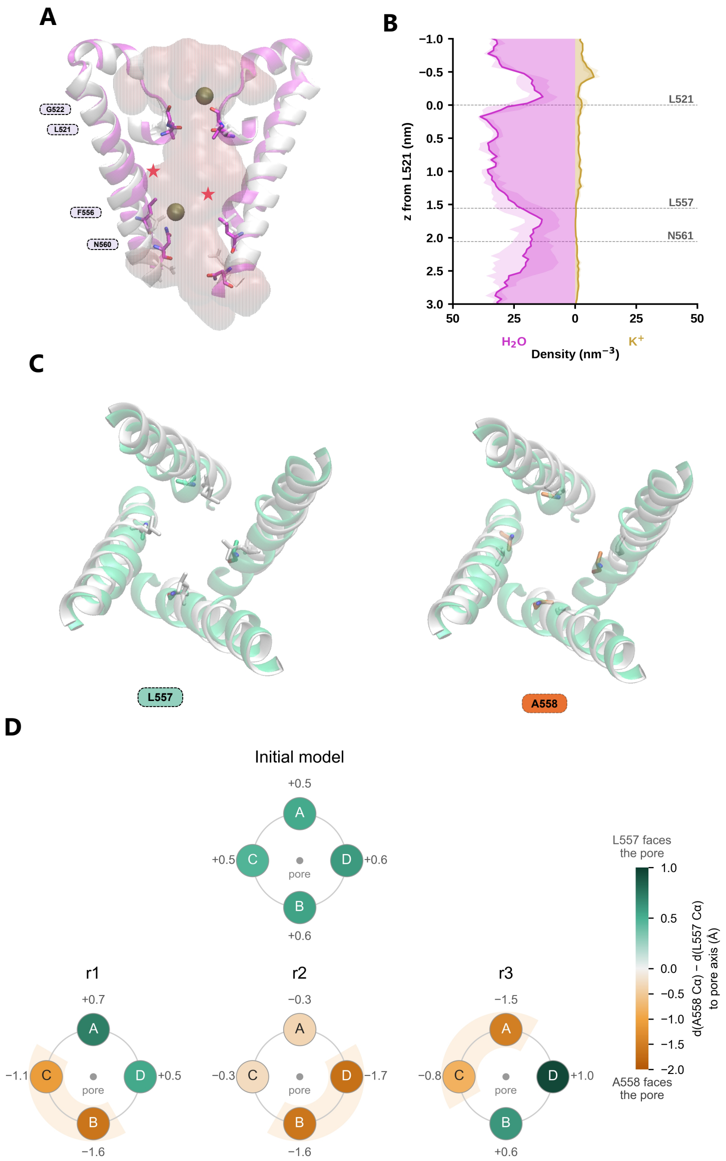


Figure S20: **Unbiased simulations of the PKD2L1 $\pi$-conformation.** (**A**) Representative snapshot of PKD2L1-$\pi$ during unbiased MD without an applied electric field, superposed on the white initial model (Fig. 5C). $\pi$-bulges are marked by red stars. Pore water is shown as a pink surface and $K^+$ ions as spheres. (**B**) Water (left) and $K^+$ (right) density within 3 Å of the pore axis, averaged over three 1-$\mu$s replicas (shading, SD across replicas). $z$ is measured from the selectivity filter (L521 C$\alpha$), with positive values towards the cytoplasm. (**C**) Top views of the pore-lining S6 helices of PKD2L1-$\pi$ in the final frame of the first of three independent 1 $\mu$s simulations (r1 in D), showing the side chains of L557 (left) and A558 (right), superposed on the white initial model (Fig. 5C). (**D**) Top-down view from the extracellular side of the initial model (top) and of three independent 1 $\mu$s simulations (r1–r3, bottom). Each circle is one subunit, drawn at 90° intervals in its counter-clockwise order around the pore. Colour and number give $\Delta = d(\text{A558 C}\alpha) - d(\text{L557 C}\alpha)$, the difference between the distances of the two C$\alpha$ atoms from the pore axis. Negative values (orange) indicate that A558 is closer to the pore axis than L557, and positive values (green) indicate that L557 is closer. For r1–r3, $\Delta$ is averaged over all frames of each 1 $\mu$s trajectory. The pore axis runs normal to the membrane through the centroid of the F480, F556 and I560 C$\alpha$ atoms. Shading marks the subunits with $\Delta < 0$ in at least 90% of frames. In the initial model, L557 faces the pore in all four subunits.

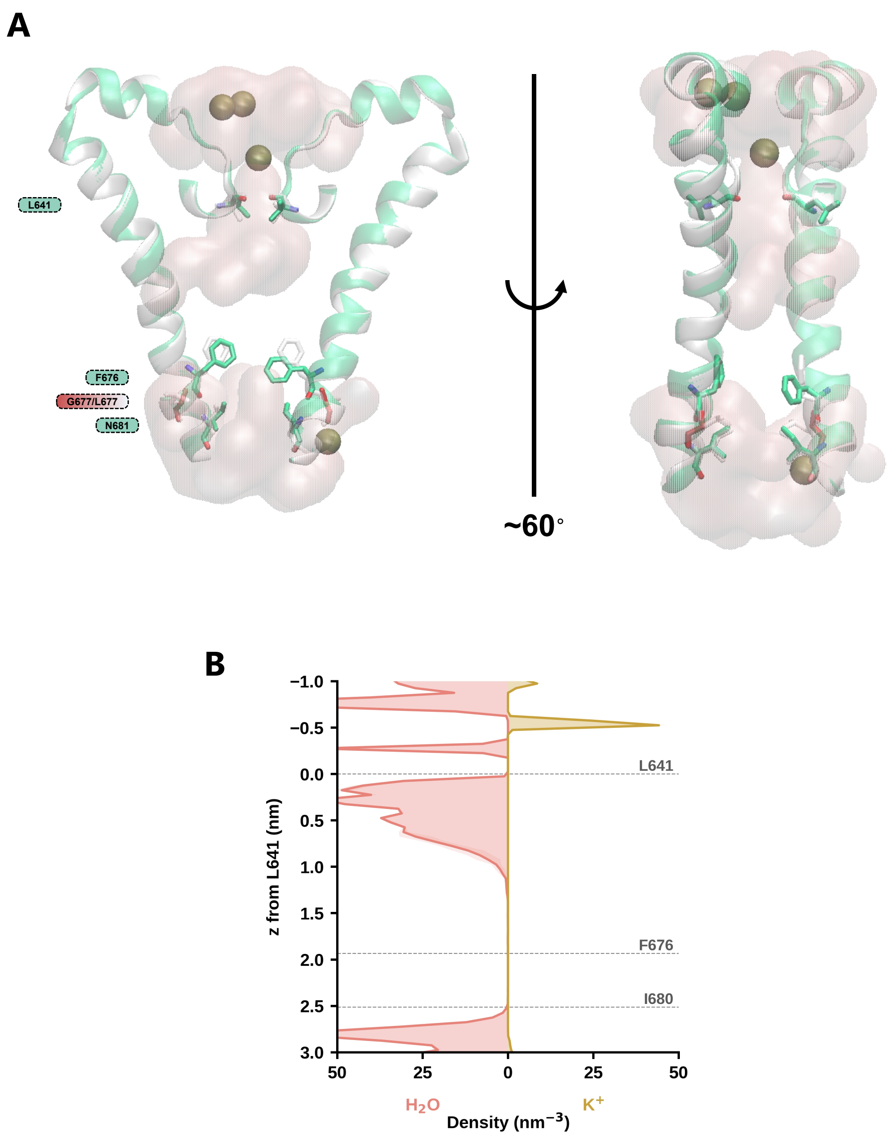


Figure S21: **L677G mutant of the $\alpha$-conformation.** (**A**) Representative snapshot of L677G with the protein backbone atoms restrained to the initial model under an applied electric field, shown in two views rotated by approximately 60° and superposed on the white initial model. Pore water is shown as a pink surface and $K^+$ ions as spheres. (**B**) Water (left) and $K^+$ (right) density within 3 Å of the pore axis, averaged over three 100-ns replicas (shading, SD across replicas). $z$ is measured from the selectivity filter (L641 C$\alpha$), with positive values towards the cytoplasm. The pore remained dewetted between F676 and I680.

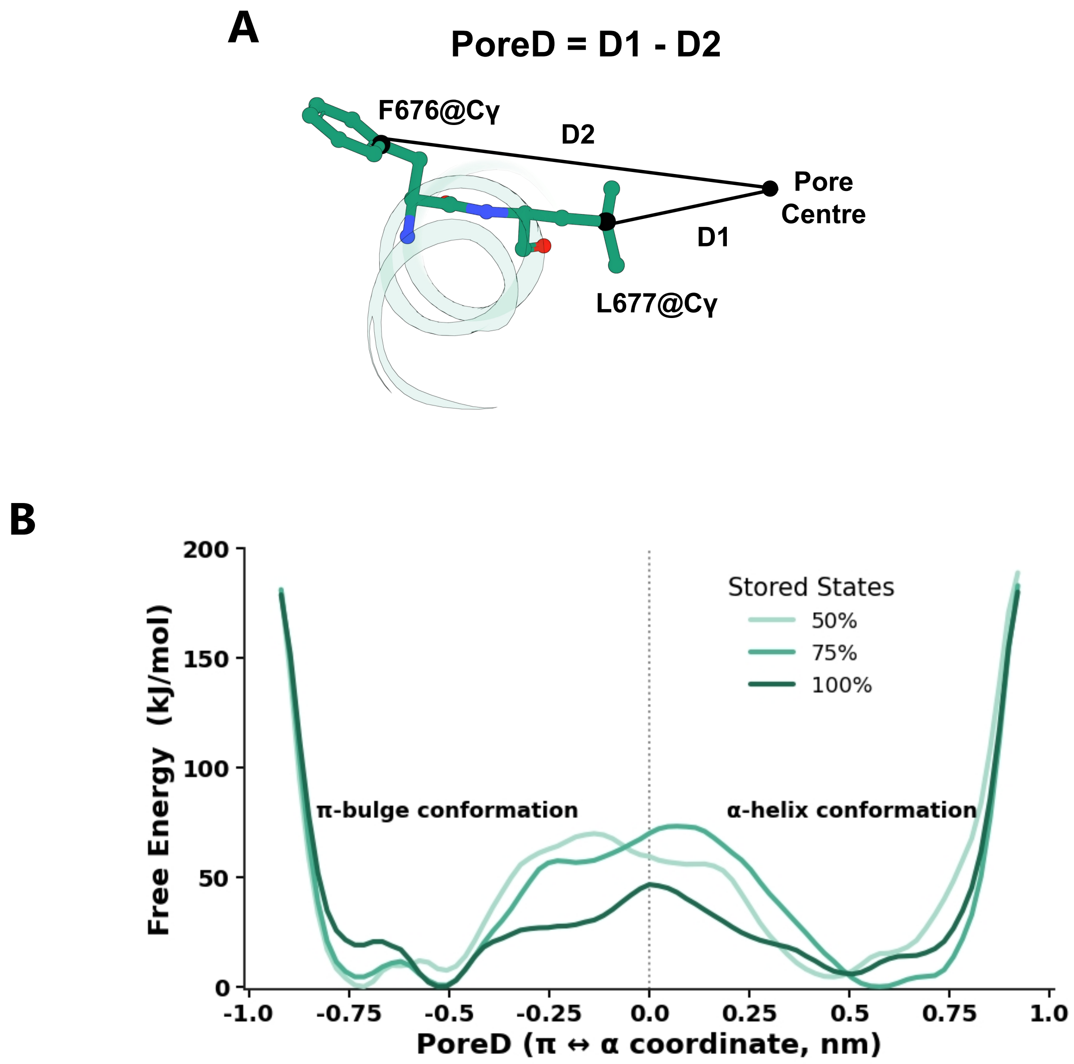


Figure S22: **Preliminary free-energy calculations of the S6 $\alpha$-to-$\pi$ transition.** (**A**) Definition of the collective variable PoreD $= D_1 - D_2$, where $D_1$ and $D_2$ are the lateral distances of the L677 and F676 C$\gamma$ atoms of one subunit from the pore axis, defined by the centroid of the V665 C$\alpha$ atoms of the four subunits. Negative values correspond to the $\pi$-bulge conformation and positive values to the $\alpha$-helical conformation. (**B**) Free-energy profile along PoreD for one subunit of WT-$\alpha$. Two-dimensional OPES-explore simulations (OPES_METAD_EXPLORE in PLUMED) were performed with PoreD and the F667 O–L672 N distance, which reports the $i \rightarrow i + 5$ hydrogen bond of the $\pi$-bulge, as collective variables. Four walkers shared a common bias, two starting from the $\alpha$-conformation and two from a $\pi$-conformation generated by steered MD, and each walker was simulated for 60 ns (240 ns in total) with CHARMM36m and TIP3P water without an applied electric field. Bias kernels were deposited every 1 ps with a barrier parameter of 100 kJ mol$^{-1}$. The free-energy surface was reconstructed from the stored OPES states using 50%, 75% and 100% of the simulation and projected onto PoreD. The profile, in particular the barrier between the two conformations, changed with the amount of data used, indicating that the calculation was not converged.

### Supplementary tables

Table S1: **Simulation systems.** All boxes had lateral dimensions of approximately 13.51 × 11.70 nm. ext., S6 extended to residue 694. Gap edit, alternative conformation built with MODELLER (see Methods). HM, homology model built from the $\pi$-conformation. $L_z$ is the box height of the initial structure with the CHARMM36m/TIP3P or AMBER ff19SB/OPC force field. — indicates that the system was not built/simulated with that set of force fields.

| | | | | $L_z$ (nm) | |
|---|---|---|---|---|---|
| System | Source | Build | Residues | CHARMM/TIP3P | AMBER/OPC |
| WT-$\alpha^{\mathrm{HM}}$ | 5T4D | Gap edit | 216–694 | 13.94 | — |
| WT-$\alpha$ | 6D1W | P604F | 216–682 | 12.89 | 13.00 |
| F604P-$\alpha$ | 6D1W | — | 216–682 | 12.66 | — |
| F676G-$\alpha$ | 6D1W | P604F, F676G | 216–682 | 12.89 | 13.00 |
| I680G-$\alpha$ | 6D1W | P604F, I680G | 216–682 | 12.89 | 13.00 |
| L677G-$\alpha$ | 6D1W | P604F, L677G | 216–682 | — | 13.00 |
| WT-$\alpha$-ext | 6D1W | P604F, ext. | 216–694 | — | 14.24 |
| F676G-$\alpha$-ext | 6D1W | P604F, F676G, ext. | 216–694 | — | 14.24 |
| I680G-$\alpha$-ext | 6D1W | P604F, I680G, ext. | 216–694 | — | 14.24 |
| WT-$\pi$ | 5T4D | — | 216–694 | 13.61 | 13.51 |
| L677G-$\pi$ | 5T4D | L677G | 216–694 | — | 13.51 |
| F676G-$\pi$ | 5T4D | F676G | 216–694 | — | 13.51 |
| PKD2L1-$\alpha$ | 5Z1W | Boltz-2 | 101–563 | — | 12.52 |
| PKD2L1-$\pi$ | 5Z1W | Boltz-2, gap edit | 101–563 | — | 12.54 |

Table S2: **Summary of production simulations.** Systems are defined in Table S1. Systems contained 0.15 M KCl unless stated. CHARMM, CHARMM36m with CHARMM-modified TIP3P water. OPC, AMBER ff19SB with OPC water and 12-6-4 ions. ECC, electronic continuum correction. SF, selectivity filter. Field, strength ($\mathrm{V\,nm^{-1}}$) and direction of the applied electric field. In the distance scans of the $\alpha$-conformation, each of the ten targets was reached by 10 ns of steered MD followed by 100 ns of production. In the ion conductance simulations of the $\pi$-conformation, the gate was opened by 10 ns of steered MD followed by 500 ns of production. Figure numbers refer to the main text unless prefixed with S. The total production simulation time was approximately 61 $\mu$s.

| System | Force field | Restraints | Field | Sampling | Figure |
|---|---|---|---|---|---|
| *PKD2 $\alpha$-conformation: unbiased and restrained simulations* | | | | | |
| WT-$\alpha^{\mathrm{HM}}$ | CHARMM/TIP3P | none | — | $3 \times 1$ $\mu$s | S1 |
| WT-$\alpha$ | CHARMM/TIP3P | none | — | $6 \times 1$ $\mu$s | 2, S2 |
| F604P-$\alpha$ | CHARMM/TIP3P | none | — | $3 \times 1$ $\mu$s | S2 |
| WT-$\alpha$ | CHARMM/TIP3P | heavy atoms | — | $1 \times 100$ ns | S3 |

*Table S2 (continued)*

| System | Force field | Restraints | Field | Sampling | Figure |
|---|---|---|---|---|---|
| WT-$\alpha$ | CHARMM/TIP3P | heavy atoms | 0.02, in | $4 \times 100$ ns | S3 |
| WT-$\alpha$ | AMBER/OPC | none | — | $1 \times 100$ ns | — |
| WT-$\alpha$ | AMBER/OPC | heavy atoms | — | $1 \times 100$ ns | — |
| WT-$\alpha$ | AMBER/OPC | heavy atoms | 0.02, in | $4 \times 100$ ns | 2 |
| *PKD2 $\alpha$-conformation: gate opening and mutations* | | | | | |
| WT-$\alpha$ | AMBER/OPC | F676 scan | 0.02, in | $10 \times (10 + 100)$ ns | 3, S5 |
| WT-$\alpha$ | AMBER/OPC | F676 scan, SF wall[a] | 0.02, in | $10 \times (10 + 100)$ ns | S6 |
| WT-$\alpha$ | AMBER/OPC | I680 scan | 0.02, in | $10 \times (10 + 100)$ ns | 3, S5 |
| WT-$\alpha$ | AMBER/OPC | I680 scan, SF wall[a] | 0.02, in | $10 \times (10 + 100)$ ns | S7 |
| F676G-$\alpha$ | AMBER/OPC | none | — | $1 \times 100$ ns | — |
| F676G-$\alpha$ | AMBER/OPC | backbone | 0.02, in | $3 \times 100$ ns | 3 |
| I680G-$\alpha$ | AMBER/OPC | none | — | $1 \times 100$ ns | — |
| I680G-$\alpha$ | AMBER/OPC | backbone | 0.02, in | $3 \times 100$ ns | 3 |
| L677G-$\alpha$ | AMBER/OPC | none | — | $1 \times 100$ ns | — |
| L677G-$\alpha$ | AMBER/OPC | backbone | 0.02, in | $3 \times 100$ ns | S21 |
| WT-$\alpha$-ext | AMBER/OPC | none | — | $1 \times 1$ $\mu$s | — |
| WT-$\alpha$-ext | AMBER/OPC | F676 scan | 0.02, in | $10 \times (10 + 100)$ ns | S9 |
| WT-$\alpha$-ext | AMBER/OPC | I680 scan | 0.02, in | $10 \times (10 + 100)$ ns | S10 |
| F676G-$\alpha$-ext | AMBER/OPC | backbone | 0.02, in | $3 \times 100$ ns | S11 |
| I680G-$\alpha$-ext | AMBER/OPC | backbone | 0.02, in | $3 \times 100$ ns | S11 |
| WT-$\alpha$ | CHARMM/TIP3P | F676 scan | 0.02, in | $10 \times (10 + 100)$ ns | S12 |
| WT-$\alpha$ | CHARMM/TIP3P | I680 scan | 0.02, in | $10 \times (10 + 100)$ ns | S13 |
| F676G-$\alpha$ | CHARMM/TIP3P | backbone | 0.02, in | $3 \times 100$ ns | S14 |
| I680G-$\alpha$ | CHARMM/TIP3P | backbone | 0.02, in | $5 \times 100$ ns | S14 |
| *PKD2 $\pi$-conformation* | | | | | |
| WT-$\pi$ | AMBER/OPC | none | — | $1 \times 100$ ns | — |
| L677G-$\pi$ | AMBER/OPC | none | — | $1 \times 100$ ns | S15 |
| L677G-$\pi$ | AMBER/OPC | backbone | 0.02, in | $3 \times 100$ ns | 4 |
| F676G-$\pi$ | AMBER/OPC | none | — | $1 \times 100$ ns | — |

*Table S2 (continued)*

| System | Force field | Restraints | Field | Sampling | Figure |
|---|---|---|---|---|---|
| F676G-$\pi$ | AMBER/OPC | backbone | 0.02, in | $1 \times 100$ ns | — |
| WT-$\pi$ | AMBER/OPC | L677 scan | 0.02, in | $1 \times 100$ ns | 4 |
| WT-$\pi$ | AMBER/OPC | L677 scan, no L677 $\chi_1$ | 0.02, in | $1 \times 100$ ns | S16 |
| WT-$\pi$ | AMBER/OPC | gate open | 0.02, in | $4 \times (10 + 500)$ ns | S17, S18 |
| WT-$\pi$ | AMBER/OPC | gate open | 0.02, out | $4 \times (10 + 500)$ ns | S17 |
| WT-$\pi$ (NaCl) | AMBER/OPC | gate open | 0.02, in | $4 \times (10 + 500)$ ns | S17 |
| WT-$\pi$ (NaCl) | AMBER/OPC | gate open | 0.02, out | $4 \times (10 + 500)$ ns | S17 |
| WT-$\pi$ ($CaCl_2$) | AMBER/OPC | gate open | 0.02, in | $4 \times (10 + 500)$ ns | S17 |
| WT-$\pi$ ($CaCl_2$) | AMBER/OPC | gate open | 0.02, out | $4 \times (10 + 500)$ ns | S17 |
| WT-$\pi$ ($CaCl_2$) | AMBER/OPC | gate open | 0.05, in | $4 \times (10 + 500)$ ns | S17 |
| WT-$\pi$ ($CaCl_2$) | AMBER/OPC | gate open | 0.05, out | $4 \times (10 + 500)$ ns | S17 |
| WT-$\pi$ | CHARMM/TIP3P, ECC 0.75 | gate open | 0.02, in | $4 \times (10 + 500)$ ns | S17 |
| WT-$\pi$ | CHARMM/TIP3P, ECC 0.78 | gate open | 0.02, in | $4 \times (10 + 500)$ ns | 4, S18 |
| WT-$\pi$ | CHARMM/TIP3P, ECC 0.78 | gate open | 0.02, out | $4 \times (10 + 500)$ ns | 4 |
| WT-$\pi$ (NaCl) | CHARMM/TIP3P, ECC 0.78 | gate open | 0.02, in | $4 \times (10 + 500)$ ns | 4 |
| WT-$\pi$ (NaCl) | CHARMM/TIP3P, ECC 0.78 | gate open | 0.02, out | $4 \times (10 + 500)$ ns | 4 |
| WT-$\pi$ ($CaCl_2$) | CHARMM/TIP3P, ECC 0.87 | gate open | 0.05, in | $4 \times (10 + 500)$ ns | 4 |
| WT-$\pi$ ($CaCl_2$) | CHARMM/TIP3P, ECC 0.87 | gate open | 0.05, out | $4 \times (10 + 500)$ ns | 4 |
| *PKD2L1* | | | | | |
| PKD2L1-$\alpha$ | AMBER/OPC | none | — | $1 \times 500$ ns | S19 |
| PKD2L1-$\alpha$ | AMBER/OPC | heavy atoms | — | $1 \times 100$ ns | — |
| PKD2L1-$\alpha$ | AMBER/OPC | heavy atoms | 0.02, in | $4 \times 100$ ns | 5 |
| PKD2L1-$\pi$ | AMBER/OPC | none | — | $3 \times 1$ $\mu$s | S20 |
| PKD2L1-$\pi$ | AMBER/OPC | heavy atoms | 0.02, in | $4 \times 100$ ns | 5 |

[a] Lower walls on the diagonal (6.0 Å) and adjacent-subunit (4.2 Å) L641 O distances (Fig. S4A).

Table S3: **Structures analysed, by channel.** 541 structures from 22 channels. Species are given with their number of structures. The S6 conformation was assigned per structure (see Methods). Per-figure omissions are listed in Table S5.

| Channel | Species (n) | $\pi$ | $\alpha$ | PDB entries, $\pi$ S6 | PDB entries, $\alpha$ S6 |
|---|---|---|---|---|---|
| **TRPA** | | | | | |
| TRPA1 | human (17) | 17 | 0 | 3J9P, 6PQO, 6PQP, 6PQQ, 6V9V, 6V9W, 6V9X, 6V9Y, 6WJ5, 6X2J, 7JUP, 7OR0, 7OR1, 9M8N, 9M8S, 9MOE, 9VU1 | |
| **TRPC** | | | | | |
| TRPC3 | human (17) | 2 | 15 | 5ZBG, 9OPU | 6CUD, 7DXB, 7DXC, 7DXD, 7DXE, 9KDB, 9KDC, 9KDD, 9KDE, 9OLK, 9OLL, 9OLM, 9OLX, 9U5C, 9VFI |
| TRPC4 | mouse (2), human (1) | 3 | 0 | 5Z96, 6JZO, 8WPN | |
| TRPC5 | human (25), mouse (2) | 26 | 1 | 6AEI, 6YSN, 7D4P, 7D4Q, 7E4T, 7WDB, 7X6C, 7X6I, 8GVW, 8GVX, 9G4Y, 9G50, 9GKB, 9GL6, 9KHJ, 9KHK, 9L3F, 9LZY, 9LZZ, 9M36, 9M4W, 9M5V, 9RRF, 9RRM, 9RRN, 9RSH | 9RRO |
| TRPC6 | human (5) | 2 | 3 | 5YX9, 7DXF | 6UZ8, 6UZA, 7DXG |
| **TRPM** | | | | | |

*continued on next page*

*Table S3 (continued)*

| Channel | Species (n) | $\pi$ | $\alpha$ | PDB entries, $\pi$ S6 | PDB entries, $\alpha$ S6 |
|---|---|---|---|---|---|
| TRPM2 | human (14) | 13 | 1 | 6MIX, 6MIZ, 6PUO, 6PUR, 6PUS, 6PUU, 7VQ1, 7VQ2, 8E6Q, 8E6R, 8E6S, 8E6T, 8E6U | 6MJ2 |
| TRPM3 | rabbit (23), mouse (14), human (1) | 38 | 0 | 8DDQ, 8DDR, 8DDS, 8DDT, 8DDU, 8DDV, 8DDX, 8ED7, 8ED8, 8ED9, 9B28, 9B29, 9B2A, 9J84, 9N1G, 9N1H, 9N1J, 9N1L, 9N1M, 9N1N, 9N4J, 9O7W, 9O7X, 9O7Y, 9O7Z, 9O80, 9O81, 9O82, 9O83, 9O86, 9O87, 9O88, 9O89, 9O8C, 9O8D, 9O8F, 9O8G, 9QHN | |

*continued on next page*

*Table S3 (continued)*

| Channel | Species (n) | $\pi$ | $\alpha$ | PDB entries, $\pi$ S6 | PDB entries, $\alpha$ S6 |
|---|---|---|---|---|---|
| TRPM4 | human (37), mouse (7) | 44 | 0 | 30KH, 30KW, 30KZ, 30LA, 30LD, 5WP6, 6BCJ, 6BCL, 6BCO, 6BCQ, 6BQR, 6BQV, 6BWI, 8RCR, 8RCU, 8RD9, 9B8W, 9B8Y, 9B90, 9B92, 9B93, 9B94, 9I3R, 9MRT, 9MT8, 9MTA, 9MTC, 9SMK, 9Y2A, 9Y2B, 9YVK, 9YVL, 9YVM, 9YVN, 9YVO, 9YVP, 9Z1W, 9Z1Z, 9Z20, 9Z21, 9Z22, 9Z23, 9Z25, 9Z27 | |
| TRPM5 | rat (8) | 8 | 0 | 8SL6, 8SL8, 8SLA, 8SLE, 8SLI, 8SLP, 8SLQ, 8SLW | |
| TRPM7 | mouse (12) | 12 | 0 | 5ZX5, 6BWD, 6BWF, 8SI2, 8SI3, 8SI4, 8SI5, 8SI6, 8SI7, 8SI8, 8SIA, 8W2L | |
| TRPM8 | mouse (27), human (6) | 18 | 15 | 7WRA, 7WRB, 7WRC, 7WRD, 7WRE, 7WRF, 8E4L, 9B6D, 9B6E, 9B6F, 9B6G, 9B6H, 9B6K, 9VJR, 9VNA, 9VNB, 9VNC, 9ZCV | 8BDC, 8E4M, 8E4N, 8E4O, 8E4P, 9B6J, 9P8Y, 9PB5, 9VJN, 9VJO, 9VJP, 9VJQ, 9VN9, 9ZCU, 9ZF0 |
| **TRPML** | | | | | |

*continued on next page*

*Table S3 (continued)*

| Channel | Species (n) | $\pi$ | $\alpha$ | PDB entries, $\pi$ S6 | PDB entries, $\alpha$ S6 |
|---|---|---|---|---|---|
| TRPML1 | human (22), mouse (9) | 30 | 1 | 5WJ5, 5WJ9, 5WPQ, 5WPT, 5WPV, 6E7P, 6E7Y, 6E7Z, 7MGL, 7SQ6, 7SQ7, 7SQ8, 7SQ9, 9CBZ, 9CC2, 9EB9, 9EBA, 9EKS, 9EKT, 9EKV, 9HJ6, 9HJ8, 9HL3, 9HL4, 9HL6, 9HL8, 9HLA, 9HLB, 9HLC, 9HLD | 9EKU |
| TRPML2 | mouse (9), human (6) | 8 | 7 | 7DYS, 9EKW, 9EKX, 9EKZ, 9EL1, 9Y5I, 9Y70, 9Y7O | 9EKY, 9EL0, 9Y5J, 9Y6P, 9Y71, 9Y73, 9Y75 |
| TRPML3 | human (3), marmoset (1) | 4 | 0 | 5W3S, 6AYE, 6AYF, 6AYG | |
| **TRPP** | | | | | |
| PKD2 | human (14) | 11 | 3 | 5K47, 5MKE, 5MKF, 5T4D, 6T9N, 6T9O, 6WB8, 9DLI, 9DWQ, 9DWT, 9NJS | 6D1W, 8HK7, 8K3S |
| PKD2L1 | mouse (1), human (1) | 0 | 2 | | 5Z1W, 6DU8 |
| **TRPV** | | | | | |

*continued on next page*

*Table S3 (continued)*

| Channel | Species (n) | $\pi$ | $\alpha$ | PDB entries, $\pi$ S6 | PDB entries, $\alpha$ S6 |
|---|---|---|---|---|---|
| TRPV1 | rat (60), human (15), squirrel (3) | 68 | 10 | 11CJ, 11CK, 11CL, 11CN, 11CO, 3J5P, 3J5Q, 3J5R, 3J9J, 5IRX, 5IRZ, 5IS0, 7L2H, 7L2I, 7L2J, 7L2K, 7L2L, 7L2M, 7L2N, 7L2O, 7L2P, 7L2R, 7L2S, 7L2V, 7L2W, 7L2X, 7LP9, 7LPA, 7LPB, 7LPC, 7LPD, 7LPE, 7LQY, 7LQZ, 7MZ6, 7MZ7, 7MZ9, 7MZA, 7MZB, 7MZC, 7MZE, 7RQU, 7RQV, 7RQW, 7RQX, 7RQY, 7RQZ, 8GF8, 8GF9, 8GFA, 8JQR, 8T0E, 8T0Y, 8T10, 8U30, 8U43, 8U4D, 8X94, 8YCP, 9ISG, 9OGK, 9P6B, 9PEA, 9VTZ, 9VU0, 9W4M, 9W4N, 9W4T | 7L2T, 7L2U, 7LR0, 7MZ5, 7MZD, 8T0C, 8T3M, 8U2Z, 8U3J, 8U3L |

*continued on next page*

*Table S3 (continued)*

| Channel | Species (n) | $\pi$ | $\alpha$ | PDB entries, $\pi$ S6 | PDB entries, $\alpha$ S6 |
|---|---|---|---|---|---|
| TRPV2 | rat (34), mouse (7), rabbit (4), human (3) | 12 | 36 | 11HZ, 11IA, 11IB, 6BO4, 6BO5, 7XEM, 7XEO, 7XER, 7XEU, 7XEV, 7XEW, 7YEP | 11ID, 11IE, 5AN8, 5HI9, 6OO3, 6OO4, 6OO5, 6U84, 6U86, 6U88, 6U8A, 6WKN, 7N0M, 7N0N, 7T37, 7T38, 7ZJD, 7ZJE, 7ZJG, 7ZJH, 7ZJI, 8EKP, 8EKQ, 8EKR, 8EKS, 8FFL, 8FFM, 8SLX, 8SLY, 9B3U, 9B3V, 9B3W, 9B3X, 9B3Y, 9B3Z, 9MKF |
| TRPV3 | human (42), mouse (18) | 40 | 20 | 13LJ, 13LK, 13LL, 13LM, 13LO, 13LQ, 13LU, 6DVZ, 6LGP, 6MHS, 6OT5, 6PVM, 6PVN, 6PVO, 6PVP, 6UW4, 6UW6, 6UW8, 7MIJ, 7MIK, 7MIL, 7MIM, 7MIN, 7MIO, 7UGG, 7XJ0, 7XJ3, 8GKA, 8V6K, 8V6L, 8V6N, 9BKU, 9JDM, 9JE5, 9JEF, 9JEG, 9PJ5, 9WV5, 9WV6, 9WV7 | 13LN, 13LP, 13LR, 6DVW, 6DVY, 6MHO, 6MHV, 6MHW, 6MHX, 6OT2, 6PVL, 6UW9, 7RAS, 7RAU, 7XJ2, 8V6M, 8V6O, 9JEE, 9PJ6, 9UED |
| TRPV4 | human (11), mouse (5) | 8 | 8 | 7AA5, 8FC8, 8FCA, 8FCB, 8J1B, 8J1H, 8T1B, 8T1E | 8FC7, 8FC9, 8J1D, 8J1F, 8JKM, 8JU6, 8JVJ, 8T1F |

*continued on next page*

*Table S3 (continued)*

| **Channel** | **Species (n)** | $\pi$ | $\alpha$ | **PDB entries, $\pi$ S6** | **PDB entries, $\alpha$ S6** |
|---|---|---|---|---|---|
| TRPV5 | rabbit (28) | 27 | 1 | 6DMR, 6DMU, 6DMW, 6O1N, 6O1P, 6O1U, 6O20, 6PBE, 6PBF, 7T6J, 7T6K, 7T6L, 7T6M, 7T6N, 7T6O, 7T6P, 7T6Q, 7T6R, 8FFN, 8FFO, 8FFQ, 8FHH, 8FHI, 8TF2, 8TF3, 8TF4, 9O6G | 6B5V |
| TRPV6 | human (25), rat (2) | 19 | 8 | 12AG, 6BO8, 6BO9, 6D7S, 6E2F, 6E2G, 7K4A, 7K4B, 7K4C, 7K4D, 7K4E, 7K4F, 7S88, 7S89, 8FOB, 9CUH, 9CUI, 9CUJ, 9CUK | 6BOA, 6BOB, 6D7T, 7S8B, 7S8C, 8FOA, 8SP8, 9NQ9 |

Table S4: **Residues measured, by channel.** X is the residue equivalent to PKD2 L677 in $\pi$- or F676 in $\alpha$-conformation. Each species keeps its own numbering. Species with different numbering are listed on separate rows. Residues were located in every chain by an S6 sequence motif and checked visually.

| | | **F676/L677 equivalent** | | **I680/N681 equivalent** | | **Structures** | |
|---|---|---|---|---|---|---|---|
| **Channel** | **Species** | $\pi$ (X) | $\alpha$ (X−1) | $\pi$ (X+4) | $\alpha$ (X+3) | $\pi$ | $\alpha$ |
| **TRPA** | | | | | | | |
| TRPA1 | human | I957 | L956 | V961 | A960 | 17 | 0 |
| **TRPC** | | | | | | | |
| TRPC3 | human | I655 | L654 | N659 | I658 | 2 | 15 |
| TRPC4 | mouse, human | I617 | L616 | N621 | M620 | 3 | 0 |
| TRPC5 | human, mouse | I621 | L620 | N625 | M624 | 26 | 1 |
| TRPC6 | human | I724 | L723 | N728 | I727 | 2 | 3 |
| **TRPM** | | | | | | | |
| TRPM2 | human | I1045 | L1044 | N1049 | F1048 | 13 | 1 |
| TRPM3 | rabbit | I1088 | L1087 | N1092 | F1091 | 23 | 0 |
| | mouse | I1111 | L1110 | N1115 | F1114 | 14 | 0 |
| | human | I1134 | L1133 | N1138 | F1137 | 1 | 0 |
| TRPM4 | human | I1040 | L1039 | S1044 | F1043 | 37 | 0 |
| | mouse | I1036 | L1035 | S1040 | F1039 | 7 | 0 |
| TRPM5 | rat | I974 | L973 | S978 | F977 | 8 | 0 |
| TRPM7 | mouse | I1093 | L1092 | N1097 | F1096 | 12 | 0 |
| TRPM8[a] | mouse, human | V976 | F979 (X+3) | G980 | V983 (X+7) | 18 | 15 |
| **TRPML** | | | | | | | |
| TRPML1 | human, mouse | I514 | F513 | T518 | I517 | 30 | 1 |
| TRPML2 | mouse, human | I503 | F502 | T507 | I506 | 8 | 7 |

*continued on next page*

*Table S4 (continued)*

| | | **F676/L677 equivalent** | | **I680/N681 equivalent** | | **Structures** | |
|---|---|---|---|---|---|---|---|
| **Channel** | **Species** | $\pi$ (X) | $\alpha$ (X−1) | $\pi$ (X+4) | $\alpha$ (X+3) | $\pi$ | $\alpha$ |
| TRPML3 | human, marmoset | I498 | F497 | T502 | I501 | 4 | 0 |
| **TRPP** | | | | | | | |
| PKD2 | human | L677 | F676 | N681 | I680 | 11 | 3 |
| PKD2L1 | mouse, human | L557 | F556 | N561 | I560 | 0 | 2 |
| **TRPV** | | | | | | | |
| TRPV1 | rat | I679 | L678 | G683 | M682 | 51 | 9 |
| | human | I680 | L679 | G684 | M683 | 15 | 0 |
| | squirrel | I681 | L680 | G685 | M684 | 2 | 1 |
| TRPV2 | rat, human | I642 | L641 | S646 | M645 | 5 | 32 |
| | mouse | I637 | L636 | S641 | M640 | 7 | 0 |
| | rabbit | I640 | L639 | S644 | M643 | 0 | 4 |
| TRPV3 | human, mouse | I674 | L673 | G678 | M677 | 40 | 20 |
| TRPV4 | human, mouse | I715 | L714 | G719 | M718 | 8 | 8 |
| TRPV5 | rabbit | I575 | F574 | G579 | M578 | 27 | 1 |
| TRPV6 | human | I575 | L574 | G579 | M578 | 18 | 7 |
| | rat | I574 | L573 | G578 | M577 | 1 | 1 |

[a] For TRPM8 $\alpha$ structures, F979 (X+3) and V983 (X+7) were used, as determined by visual inspection.

Table S5: **Structures excluded from the lower-gate survey.** 143 of 684 TRP EM entries in the PDB (retrieved August 2026) were removed by the general filters, leaving 541. At each position, structures in which the measured residue faces away from the pore were additionally omitted (C$\alpha$ →C$\beta$ vector more than 90° from the direction to the pore axis, or side chain reaching less than 1.0 Å beyond its C$\alpha$ towards the axis), as confirmed by visual inspection.

| Scope | Reason | n | PDB entries |
|---|---|---|---|
| Both positions | Not mammalian | 76 | 5VKQ, 6BBJ, 6BPQ, 6D73, 6DRJ, 6DRK, 6G1K, 6NR2, 6NR3, 6NR4, 6O6A, 6O6R, 6O72, 6O77, 6PKV, 6PKW, 6PKX, 6WHG, 7B05, 7B0J, 7B0S, 7B16, 7B1G, 7MBP, 7MBQ, 7MBR, 7MBS, 7MBT, 7MBU, 7MBV, 7YKR, 7YKS, 8E4Q, 8SR7, 8SR8, 8SR9, 8SRA, 8SRB, 8SRC, 8SRD, 8SRE, 8SRF, 8SRG, 8SRH, 8SRI, 8SRJ, 8SRK, 9B6I, 9FXL, 9FXM, 9JJE, 9P3N, 9P3O, 9P3P, 9P3Q, 9P3R, 9P3S, 9P3T, 9P3U, 9P3V, 9P3W, 9P7S, 9P90, 9P91, 9PAR, 9PB6, 9VQQ, 9VQW, 9VQX, 9VQY, 9ZCN, 9ZCO, 9ZCP, 9ZCQ, 9ZCR, 9ZEZ |
| Both positions | Not a tetramer | 25 | 6DJR, 8E6V, 8GKG, 8SIB, 8SWN, 8T1C, 8U3A, 8U3C, 9B8X, 9B8Z, 9B91, 9DIJ, 9IQY, 9JJF, 9QHM, 9QHO, 9QHP, 9QHQ, 9T9U, 9V6J, 9Y2C, 9Z1X, 9Z1Y, 9Z24, 9Z26 |
| Both positions | Heteromer | 18 | 6A70, 7D7E, 7D7F, 8WPL, 8WPM, 8Z6B, 8Z6F, 8Z6H, 8ZKH, 8ZKR, 8ZKS, 8ZKT, 8ZKU, 8ZL8, 9K4I, 9KHI, 9L0M, 9L0W |

*continued on next page*

*Table S5 (continued)*

| **Scope** | **Reason** | **n** | **PDB entries** |
|---|---|---|---|
| Both positions | No S6 modelled | 10 | 5YDZ, 5YE1, 5YE2, 5YE5, 6CV9, 6D7L, 6DJS, 6PVQ, 7A6U, 8DDW |
| Both positions | Mixed $\alpha/\pi$ S6 among subunits | 8 | 6OO7, 8JU5, 8T3L, 9IQX, 9RRQ, 9RRU, 9RSG, 9RVV |
| Both positions | Asymmetric subunits (diagonal C$\alpha$–C$\alpha$ differ > 2 Å) | 5 | 13LS, 13LT, 7XJ1, 8JVI, 8T1D |
| Both positions | No lower-gate reference for this channel (TRPM1) | 1 | 9OR5 |
| F676/L677 equivalent | Residue does not face the pore | 3 | 8J1D, 8J1F, 8JKM |
| I680/N681 equivalent | Residue does not face the pore | 6 | 6MJ2, 8BDC, 8E4P, 9P8Y, 9VJN, 9ZCU |
